# A Polybasic Domain in aPKC Mediates Par6-Dependent Control of Membrane Targeting and Kinase Activity

**DOI:** 10.1101/588624

**Authors:** Wei Dong, Juan Lu, Xuejing Zhang, Yan Wu, Kaela Lettieri, Gerald R. Hammond, Yang Hong

## Abstract

Mechanisms coupling the atypical PKC (aPKC) kinase activity to its subcellular localization are essential for cell polarization. Unlike other members of the PKC family, aPKC has no well-defined plasma membrane (PM) or calcium binding domains, leading to the assumption that its subcellular localization relies exclusively on protein-protein interactions. Here we show that in both *Drosophila* and mammalian cells the pseudosubstrate region (PSr) of aPKC acts as a polybasic domain capable of targeting aPKC to the PM via electrostatic binding to PM PI4P and PI(4,5)P_2_. However, physical interaction between aPKC and Par-6 is required for the PM-targeting of aPKC, likely by allosterically exposing the PSr to bind PM. Binding of Par-6 also inhibits aPKC kinase activity and such inhibition can be relieved through Par-6 interaction with apical polarity protein Crumbs. Our data suggest a potential mechanism in which allosteric regulation of polybasic PSr by Par-6 couples the control of both aPKC subcellular localization and spatial activation of its kinase activity.

**eTOC Summary:** Dong et al. discover that the pseudo-substrate region (PSr) in aPKC is a polybasic domain capable of electrostatically targeting aPKC to plasma membrane. Allosteric regulation of PSr by Par-6 couples the control of both aPKC subcellular localization and spatial activation of kinase activity.

## INTRODUCTION

Polarity proteins play conserved and essential roles in regulation cell polarity and for the majority of them achieving polarized plasma membrane (PM)/cortical localization is essential for their functions (Hong, 2018; Rodriguez-Boulan and Macara, 2014). Mechanisms mediating the PM/cortical association of these polarity proteins had often been assumed to be primarily based on the intricate protein-protein interactions among polarity proteins and their regulators (Rodriguez-Boulan and Macara, 2014). Recent studies however discovered that several polarity proteins such as Lgl, Numb and Miranda contain so-called polybasic (i.e. basic-hydrophobic) domains (Bailey and Prehoda, 2015; Dong et al., 2015), which are short but highly positively charged due to their enrichment of basic Arg/Lys residues. Polybasic domains can specifically bind to PM, as the inner surface of PM is the most negatively charged membrane surface due to membrane phosphatidylserine (Yeung et al., 2008) and the unique enrichment of polyphosphoinositides PI4P and PI(4,5)P_2_ (PIP2) (Hammond et al., 2012). Moreover, polybasic domains in Lgl, Numb and Miranda also contain conserved serine residues that can be phosphorylated by another key polarity protein atypical PKC (aPKC) (Bailey and Prehoda, 2015; Dong et al., 2015). Similar to the phosphorylation of the polybasic ED domain in MARCKS by PKC (Arbuzova et al., 2002), aPKC-phosphorylation neutralizes the positive charges in a polybasic domain therefore inhibits its electrostatic binding to the PM (Bailey and Prehoda, 2015; Dong et al., 2015). Such aPKC-dependent inhibition serves an elegant mechanism to polarize the PM targeting of polybasic polarity proteins, allowing apically localized aPKC to limit Lgl, Numb and Miranda to basolateral PM. Potential aPKC-phosphorylatable polybasic domains have been found in hundreds of proteins in metazoan genomes (Bailey and Prehoda, 2015)(Y.H. unpublished data), suggesting aPKC plays a critical role in regulating the PM targeting of many polybasic proteins.

However, to date the exact molecular mechanism governing the PM/cortical localization of aPKC itself remains unclear. Unlike conventional PKC (cPKC) and novel PKC (nPKC) that bind DAG (diacylglycerol), phospholipids and calcium for their kinase activation and PM targeting, aPKC has no well-defined PM or calcium binding domains and has not been demonstrated or proposed to directly bind the PM (Garg et al., 2013; Rosse et al., 2010). In fact, it is considered a unique feature of aPKC that its kinase activity and subcellular localizations appear to be exclusively regulated by protein-protein interactions other than lipids and/or calcium (Rosse et al., 2010). aPKC has a PB1 domain that binds to the PB1 domain in another polarity protein Par-6 which consistently colocalizes with aPKC in many polarized cell types including epithelial cells and *C. elegans* one-cell embryos (Hong, 2018; Suzuki and Ohno, 2006). PM/cortical localization of aPKC/Par-6 complex has been assumed mainly based on protein interactions with other polarity proteins such as Baz (Izumi et al., 1998; Krahn et al., 2010; Morais-de-Sa et al., 2010), Crb (Sotillos et al., 2004), Sdt/Pals1 (Wang et al., 2004), Patj (Hurd et al., 2003) or Cdc42 (Joberty et al., 2000; Lin et al., 2000; Qiu et al., 2000). Recent studies have delineated some exciting detail mechanisms by which Par-3 and Cdc42 coordinate the spatial and temprol control of aPKC kinase activity during the anterior-posterior (A-P) polarization of worm one-cell embryo (Dickinson et al., 2017; Rodriguez et al., 2017), but how aPKC PM localization and kinase activity are regulated during apical-basal (A-B) polarization is much less clear. Moreover, in vitro kinase assays yielded conflicting results regarding whether binding of Par-6 to aPKC inhibits or activates its kinase activity (Hong, 2018).

Here we report that the pseudosubstrate region (**PSr**) of aPKC also functions as a polybasic domain that directly binds to PM through electrostatic interaction with PM phosphoinositides PI4P and PIP2. Interestingly, unlike the polybasic domains in Lgl, Numb and Miranda, PSr in aPKC has not been shown to be phosphorylatable. Instead we show that the PM binding of PSr appears to be allosterically controlled by protein interactions between aPKC and Par-6. Our data suggest a model that Par-6-dependent allosteric regulation of polybasic PSr in aPKC couples the PM targeting of aPKC and the spatially restricted activation of aPKC kinase during apical-basal polarization.

## RESULTS

### The conserved pseudosubstrate region (PSr) is a polybasic domain required for the PM targeting of DaPKC in *Drosophila* epithelial cells

Our previous stuides showed that acute hypoxia induces loss of PM phosphoinositides which in turn disrupts the PM localization of polybasic domain proteins such as Lgl and Numb in live fly tissues and in cultured cells (Dong et al., 2015). Curiously, in the same studies we found that endogenous *Drosophila* aPKC (DaPKC) also showed dramatic loss of PM localization in epithelial cells under hypoxia (Dong et al., 2015), suggesting that PM targeting of DaPKC may be based on direct binding to PM phospholipids. Similar to conventional PKC (cPKC) and novel PKC (nPKC), aPKC contains a pseudosubstrate region (PSr) that binds the kinase domain to induce autoinhibition (Rosse et al., 2010). We identified that the PSr in DaPKC is in fact highly polybasic (i.e. basic-hydrophobic) (Fig. 1A): the 17aa PSr contains eight Arg and Lys residues plus one Leu and one Trp residues, and the adjacent 13aa sequence in C1 domain contains additional four Arg and Lys residues plus two Phe residues (Heo et al., 2006). Besides the enriched Arg and Lys residues, the presence of Trp, Phe and Leu is highly indicative for a polybasic domain as the bulky or long hydrophobic side chains of these residues enhance the PM binding (Heo et al., 2006; McLaughlin and Murray, 2005; Yeung et al., 2006). The PSr is extremely well conserved between DaPKC and mammalian aPKC isoforms PKCζ and PKCι (Fig. 1A), with a basic-hydrophobic index (Brzeska et al., 2010) of 0.92, comparable to polybasic domains in Lgl (1.00) and Numb (0.89).

**Figure 1.**
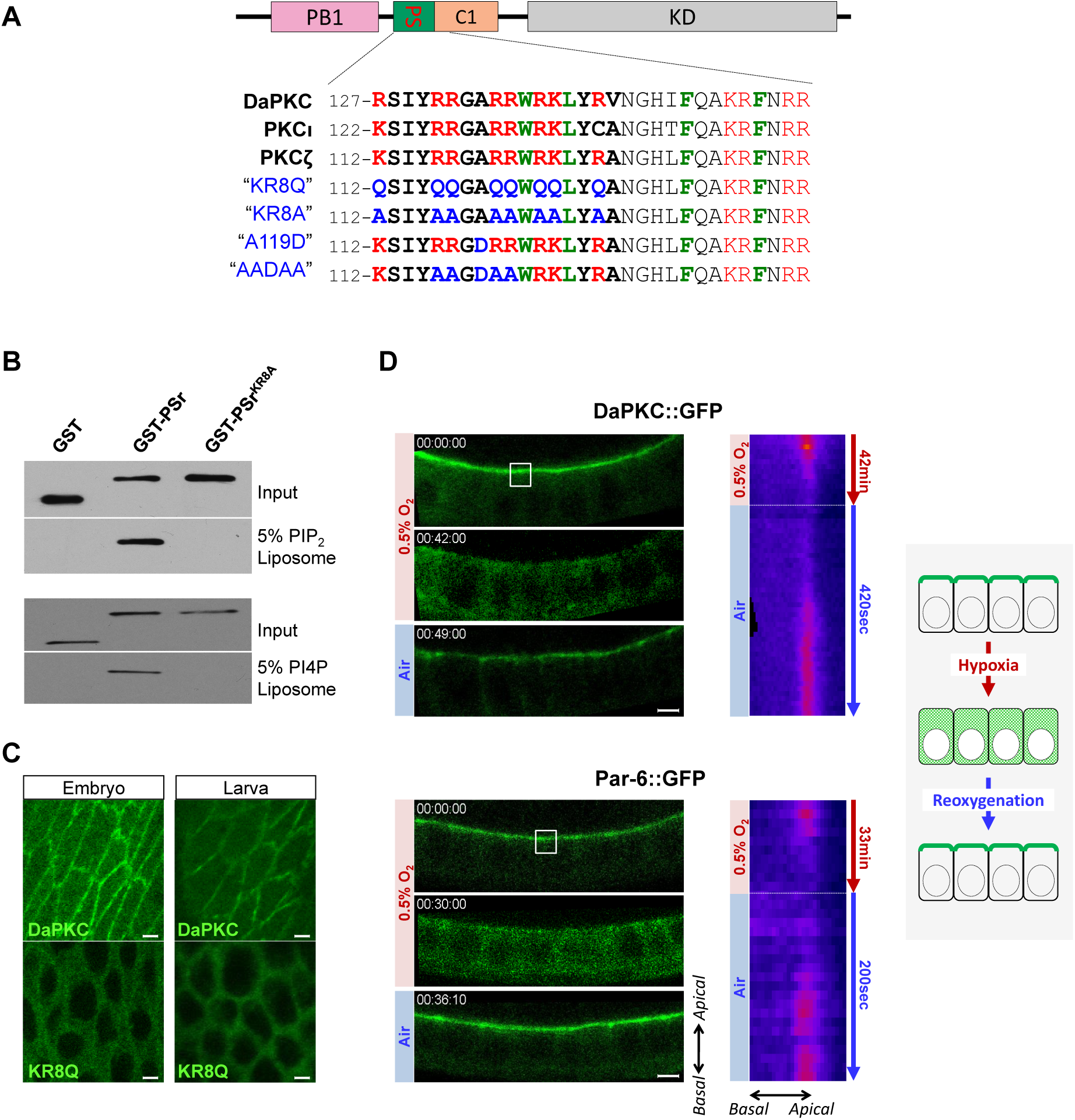
The conserved polybasic pseudosubstrate region (PSr) mediates PM targeting of aPKC in *Drosophila* epithelia. **(A)** Alignment of the PSr (bold) and adjacent sequences in C1 domain from *Drosophila* and mammalian aPKC isoforms. Sequences are based on NCBI RefeSeq# NP_524892.2 (DaPKC), NP_002735.3 (PKCζ), and NP_002731.4 (PKCι), respectively. Residues mutated in aPKC^KR8Q^ (“KR8Q”), aPKC^KR8A^ (“KR8A”), PKCζ^A119D^(“A119D”) and PKCζ^AADAA^ (“AADAA”) are also shown. **(B)** GST fusion of PSr from PKCζ (“GST-PSr”), but not the non-polybasic GST-PSr-KR8A, co-sedimented with PI4P- and PIP2-containing liposomes in vitro. **(C)** DaPKC::GFP (“DaPKC”), but not non-polybasic DaPKC^KR8Q^::GFP (“KR8Q”), localized to PM in embryonic and larval disc epithelia. **(D)** Follicular epithelial cells in ovaries from *ubi-DaPKC::GFP* or *par-6::GFP* adult females were imaged ex vivo under controlled oxygen environment. Cells are in cross-section view as indicated by the illustration at the far right. PM localization of DaPKC::GFP and Par-6::GFP were acutely inhibited by hypoxia (0.5% O_2_), but recovered after reoxygenation by air (see also Movies S1 and S2). Kymographs on the right show the acute loss and recovery DaPKC::GFP or Par-6::GFP on PM during hypoxia and post-hypoxia reoxygenation treatments. White boxes indicate where kymographs were sampled. Scale bars: 5µm (C,D).

Similar to the polybasic domain in Lgl (Dong et al., 2015), PSr in aPKC is also capable of direct binding to PI4P and PIP_2_ *in vitro* in liposome pull-down assays. Purified GST fusion of PSr from PKCζ bound PI4P- and PIP_2_-containing liposomes, while non-polybasic GST-PSr-KR8Q with all eight Arg/Lys residues in PSr mutated to Gln, did not (Fig. 1B). PSr is also required for PM targeting of DaPKC *in vivo*: while ubiquitously expressed aPKC::GFP localized to apical PM in both wild type and *aPKC*^-/-^ epithelial cells, non-polybasic aPKC^KR8Q^::GFP stayed in the cytosol (Fig. 1C and Fig. S1A).

In vivo, DaPKC::GFP also showed acute and reversible loss from PM similar to Lgl and Numb in live *Drosophila* follicular and embryonic epithelial cells under hypoxia (Fig. 1D, Fig. S1B and Movie S1), consistent with that electrostatic binding to phosphoinositides could be the primary mechanism targeting DaPKC to the PM. Furthermore, Par-6 is an essential regulatory partner of aPKC and forms a robust complex with aPKC (Hong, 2018; Suzuki and Ohno, 2006). Both proteins are mutually dependent on each other for proper localization and function during cell polarization processes. Consistently, hypoxia in live *Drosophila* follicular and embryonic epithelial cells also induced acute and reversible loss of PM targeting of Par-6::GFP (Fig. 1D, Fig. S1B and Movie S2). The loss of DaPKC/Par-6 complex from PM under hypoxia is not due to the concurrent loss of Lgl from PM. Par-6::GFP in *lgl*^-/-^ mutant follicular cells showed expanded localization to basolateral PM but responded to hypoxia identically to Par-6::GFP in wild type cells (Fig. S1C).

Overall, our data suggest that the polybasic PSr in DaPKC is essential for localizing DaPKC/Par-6 complex to the PM in *Drosophila* epithelia in vivo, likely through direct interaction with PM phosphoinositides such as PI4P and PIP2 (see below).

### PM targeting of mammalian PKCζ depends on both PSr and Par-6

Given the well documented requirement of Par-6 in aPKC subcellular localization in many cell types (Chen et al., 2018; Hong, 2018; Suzuki and Ohno, 2006), we further investigated the role of Par-6 in regulating the polybasic PSr-mediated PM targeting of aPKC. For such purpose we took the over-expression approach in non-polarized HEK293 cells to investigate the potential interactions between aPKC and Par-6 or other exogenously introduced proteins. To keep results consistent, all cultured cell experiments reported here only used mammalian proteins of Lgl, aPKC, Par-6, Crb and Cdc42. Mammalian aPKC family has two isoforms, PKCζ and PKCι, and both contain PSr domains nearly identical to DaPKC’s (Fig. 1A). A PKCζ-specific antibody (Stross et al., 2009) showed that HEK293 cells express no detectable endogenous PKCζ (Fig. 2A,B) thus provide a relatively clean background for investigating the PM targeting of the exogenously expressed PKCζ. Surprisingly, we found that PKCζ::GFP was completely cytosolic when expressed alone (Fig. 2A). However, although Par-6::RFP was also cytosolic when expressed alone, co-expression of PKCζ::GFP and Par-6::RFP resulted in strong and robust PM localization of both proteins (Fig. 2A). We also made a bicistronic construct that expresses a fusion protein of PKCζ::RFP and Par-6::iRFP connected by a 2A peptide which self-cleaves during translation (Chan et al., 2011) to produce PKCζ (“PKCζ::RFP^-2A^”) and Par-6 (“^2A-^Par-6::iRFP”) at a constant 1:1 ratio. The fusion protein appeared to be efficiently cleaved (Fig. 2C) and both PKCζ::RFP^-2A^ and ^2A-^Par-6::iRFP showed strong PM localization as expected (Fig. 2A).

**Figure 2.**
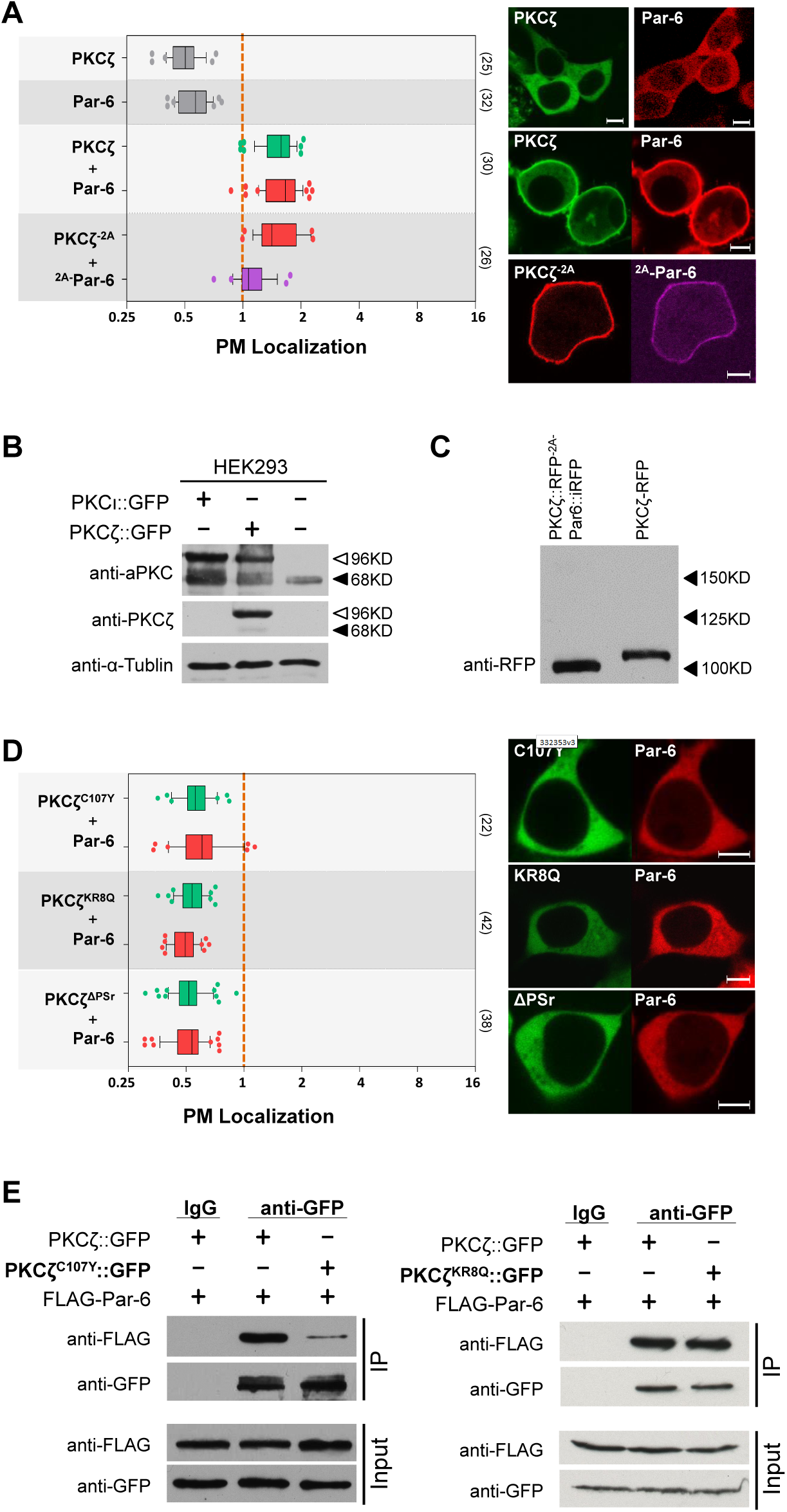
PM localization of PKCζ in HEK293 cells requires both polybasic PSr and Par-6. **(A)** PKCζ::GFP or Par-6::RFP was cytosolic when expressed alone, but both became strongly PM localized when co-expressed. PKCζ::RFP^-2A^ and ^2A-^Par-6::iRFP also showed strong PM localization. **(B)** Pan-aPKC antibody (“anti-aPKC”) detects both exogenously expressed PKCɩ::GFP and PKCζ::GFP in HEK293 cells (white arrowhead), as well as endogenous expressed aPKC (black arrowhead). PKCζ-specific antibody (“anti-PKCζ”, C24E6 from Cell Signaling) specifically detected exogenously expressed PKCζ::GFP but showed no detectable expression of endogenous PKCζ in HEK293. α-Tublin serves as loading control. **(C)** Only PKCζ::RFP^-2A^ (∼100KD) but not full-length PKCζ::RFP^-2A-^Par-6::iRFP fusion protein (∼150KD) was detected in cells expressing PKCζ::RFP^-2A-^Par-6::iRFP. Lysate from cells expressing PKCζ::RFP was loaded and blotted as a positive control. **(D)** PKCζ^C107Y^::GFP (“C107Y”), PKCζ^KR8Q^::GFP (“KR8Q”), PKCζ^ΔPSr^::GFP (“ΔPSr”) did not localize to PM when co-expressed with Par-6::RFP. **(E)** Both PKCζ::GFP and PKCζ^KR8A^::GFP, but not PKCζ^C107Y^::GFP, co-immunoprecipitated with FLAG-Par-6 by anti-GFP antibody from HEK293 cells. In all data plots, boxes extend from 25 and 75 percentiles, with lines in the middle indicating the median and whiskers indicating 10 and 90 percentiles, respectively. Sample numbers are indicated in parentheses at the right. Orange dashed lines in quantification figures indicate the PM localization index = 1 (see Materials and Methods). Measurements less than 1 indicate cytosolic localization, whereas larger than 1 indicate PM localization. PM localization axes in all figures are in log2 scale. Scale bars: 5µm (A,D).

We confirmed that such co-localization to PM requires direct physical interaction between Par-6 and PKCζ, as a C107Y mutant of PKCζ which specifically abolishes the physical interaction with Par-6 (Kim et al., 2009), failed to localize to PM when co-expressed with Par-6 (Fig. 2D,E). Furthermore, PM targeting of PKCζ/Par-6 complex requires the polybasic PSr in PKCζ. Co-expression of Par-6 with non-polybasic PKCζ^KR8Q^ resulted in no PM localization of either protein (Fig. 2D), even though the physical interaction between Par-6 and PKCζ^KR8Q^ remained intact (Fig. 2E). The loss of PM targeting of PKCζ^KR8Q^ is not due to potential misfolding, as PKCζ^KR8Q^ phosphorylates Lgl similarly to PKCζ in HEK293 cells (Fig. 6B,C). Our findings are not unique to HEK293 cells, as COS7 and MCF7 breast cancer cells also showed same Par-6-dependent PM targeting of PKCζ (Fig. S2A,B). In fully polarized MDCK cells, overexpressed PKCζ alone was cytosolic although Par-6 alone was partially PM localized (Fig. S2C). Nonetheless, co-expression of PKCζ and Par-6 resulted in robust PM localization of both proteins (Fig. S2C). Interestingly, Par-6 became cytosolic when co-expressed with PKCζ^KR8Q^ (Fig. S2C), which could indicate a dominant-negative effect of PKCζ^KR8Q^ due to the formation of PKCζ^KR8Q^/Par-6 that is incapable of PM targeting. Thus, in polarized MDCK cells additional mechanisms exist to at least partially target Par-6 to PM, but PM targeting of PKCζ remains dependent on both polybasic PSr and Par-6.

### PM targeting of aPKC in HEK293 cells is Cdc42-independent

Cdc42 has been shown to be essential for aPKC/Par-6’s localization to anterior PM in *C. elegans* one-cell embryo and has well characterized physical interactions with Par-6. We thus investigated whether aPKC/Par-6 PM targeting in HEK293 cells is also Cdc42-dependent. In HEK293 cells expressing constitutively active Cdc42^CA^, Par-6 was readily recruited to the PM as expected (Fig. 3A). Furthermore, non-polybasic PKCζ^KR8Q^ was cytosolic in cells expressing either Cdc42^CA^ (Fig. 3A) or Par-6 (Fig. 2D), but became PM-localized in cells co-expressing both Cdc42^CA^ and Par-6 (Fig. 3A). Thus, in HEK293 cells the interaction between Cdc42^CA^ and Par-6 is sufficient to recruit Par-6/aPKC complex to the PM and such PM recruitment can be independent of the polybasic PSr in aPKC. However, the role of Cdc42^CA^ in targeting aPKC/Par-6 is likely context-dependent, as in *Drosophila* follicular cells Cdc42^CA^ overexpression failed to recruit DaPKC^KR8Q^::GFP to the PM (Fig. 3B).

**Figure 3.**
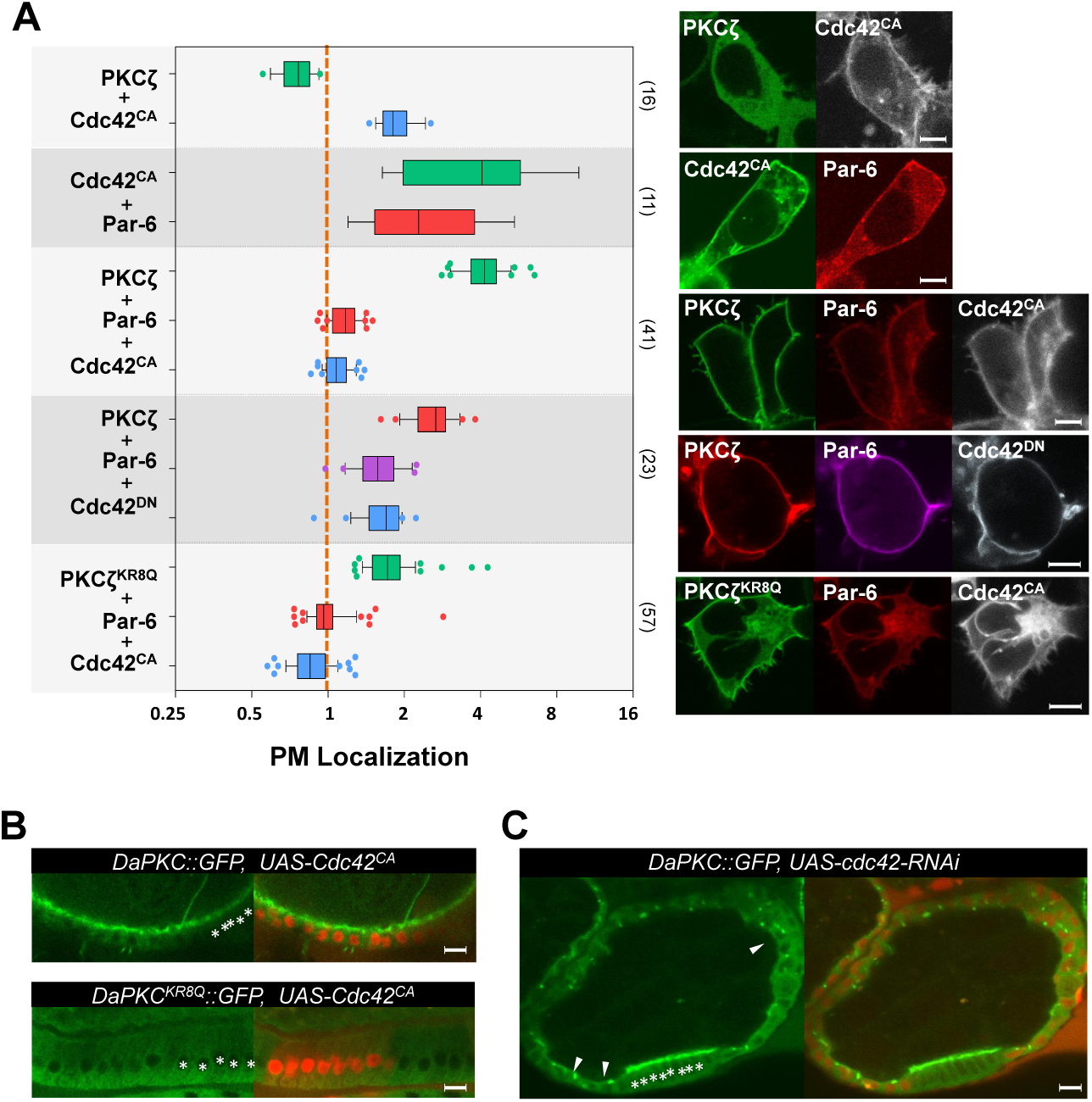
PM targeting of PKCζ in HEK293 cells is independent of Cdc42. **(A)** PKCζ::GFP was cytosolic when co-expressed with BFP::Cdc42^CA^. Par-6::RFP localized to PM in cells co-expressing GFP::Cdc42^CA^. Co-expressed PKCζ and Pra-6 were PM localized in cells expressing either BFP::Cdc42^CA^ or BFP::Cdc42^DN^. Non-polybasic PKCζ^KR8Q^::GFP was PM-localized in cells expressing both BFP::Cdc42^CA^ and Par-6::RFP. **(B)** Overexpression of Cdc42^CA^ did not change DaPKC::GFP and DaPKC^KR8Q^::GFP subcellular localization in *Drosophila* follicular epithelial cells. **(C)** DaPKC::GFP formed intracellular puncta (arrowheads) in *cdc42-RNAi* follicular cells. In B and C: cells expressing *UAS-cdc42-RNAi* or *UAS-Cdc42^CA^* are marked by RFP expression. Asterisks highlight the wild type cells. Scale bars: 5µm (A, B), 15µm (C)

On the other hand, in HEK293 cells overexpression of dominant-negative Cdc42^DN^ did not inhibit the PM targeting of aPKC/Par-6 (Fig. 3A). In *Drosophila* follicular cells subject to *cdc42-RNAi*, DaPKC::GFP was reduced from PM and enriched at intracellular puncta (Fig. 3C), likely due to the interaction between aPKC/Par-6 and polarity protein Baz which was also enriched at the same puncta (Fletcher et al., 2012). To further clarify the role of Cdc42 in our cell-based assays, we made two Par-6 mutants: PB1^Par-6^ which removes the CRIB and PDZ domains thus retains only the N-terminal PB1 domain, and Par-6^ΔPB1^ which carries the reciprocal truncation. PB1^Par-6^ was fully capable of targeting PKCζ to PM but Par-6^ΔPB1^ was not (Fig. 5B), suggesting the physical interaction between the N-terminal PB1 domains in aPKC and Par-6 is sufficient to induce PSr-dependent PM targeting of aPKC. Given that PB1^Par-6^ does not contain any known Cdc42 interacting domain, PM targeting of PKCζ by PB1^Par-6^ should be Cdc42-independent.

In summary, our results do not exclude the role of Cdc42 in regulating aPKC/Par-6 PM targeting under certain cellular contexts, but show that PM targeting of aPKC/Par-6 can be mechanistically independent of Cdc42.

### PI4P and PIP2 act redundantly to target aPKC/Par-6 complex to PM

Phosphoinositides PI4P and PIP2 are uniquely enriched in PM and are considered the major negatively charged phospholipids responsible for electrostatically binding polybasic domains (Hammond, 2012). To confirm that PM targeting of aPKC/Par-6 complex in cells is indeed mediated by PI4P and PIP2, we used a well-established system to acutely and selectively deplete PI4P, PIP2 or both in HEK293. In this inducible system addition of rapamycin induces dimerization between FKBP-tagged phosphatases and FRB-tagged PM anchor protein Lyn11-FRB-CFP, resulting in acute PM recruitment of phosphatase and concurrent depletion of target phospholipids (Hammond, 2012). In particular, PM-recruitment of a chimeric lipid phosphatase PJ (pseudojanin) rapidly converts both PI4P and PIP2 in PM to phosphatidylinositol (PI) (Hammond, 2012) (Fig. 4A). In HEK293 cells expressing PKCζ::GFP, Par-6::iRFP together with RFP-FKPB-PJ and Lyn11-FRB-CFP, addition of rapamycin induced acute PM localization of PJ and concurrent loss of both PKCζ::GFP and Par-6::iRFP from PM (Fig. 4B). Depleting either PIP2 using FKBP-INPP5E or PI4P using FKBP-PJ-Sac induced much milder loss of PM targeting of both proteins (Fig. 4A,B).

**Figure 4.**
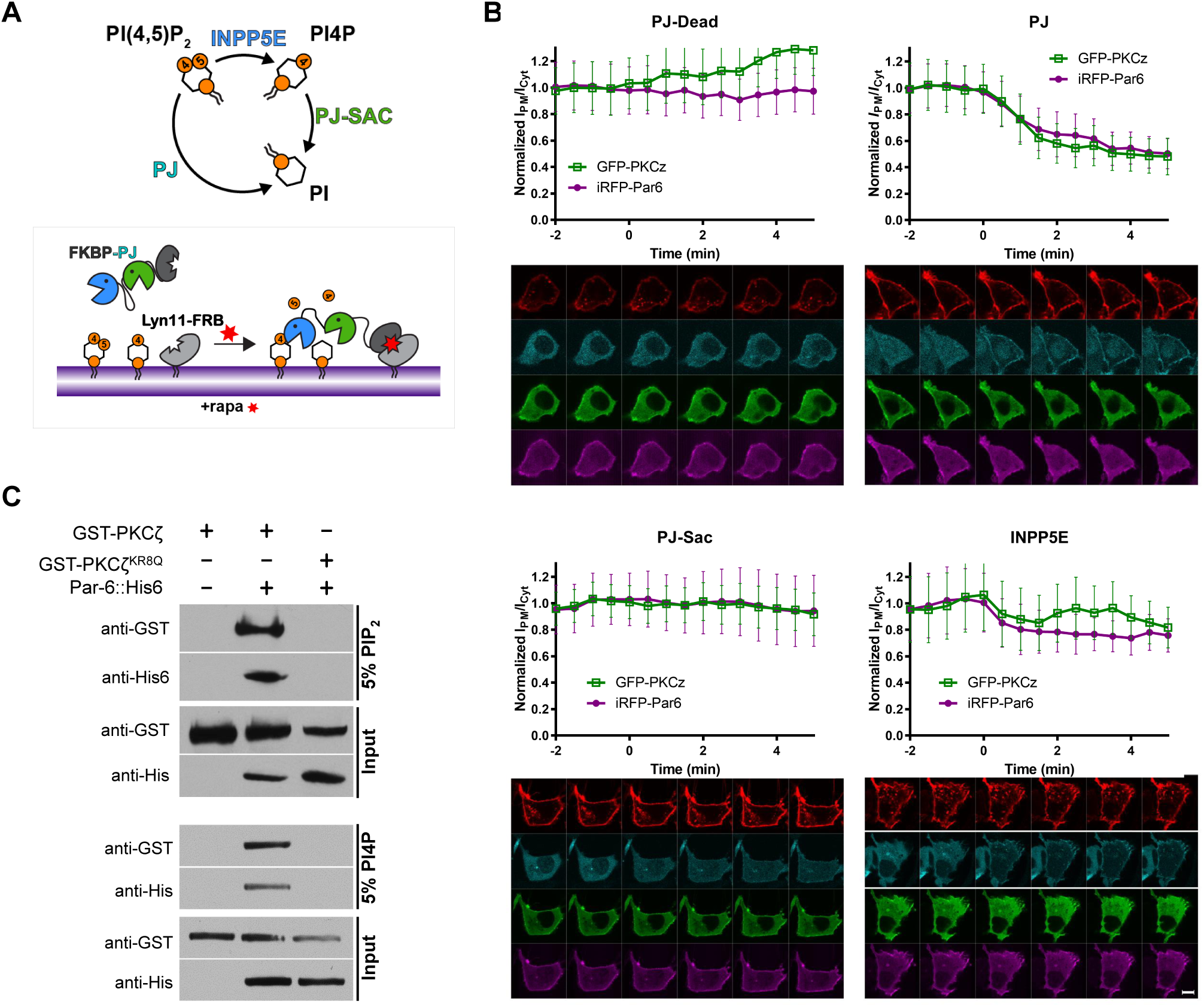
PM targeting of PKCζ and Par-6 depends on both PI4P and PIP2. **(A)** INPP5E converts PIP2 to PI4P which can be further converted to PI by Sac, whereas PJ converts both PIP2 and PI4P to PI. Box: FKBP-PJ can be acutely recruited to PM through rapamycin (rapa)-induced heterodimmerization with PM-anchored Lyn11-FRB. PM-recruitment of PJ results in acute depletion of both PI4P and PIP2. **(B)** PM localization of PKCζ::GFP and Par-6::iRFP were quantified prior and after the rapamycin addition in HEK293 cells expressing Lyn11-FRB-CFP and mCherry-FKBP-PJ or -Sac, or - INPP5E, or -PJ-dead (as a negative control). Representative time-lapse images of Lyn11-FRB- CFP (red), PKCζ::GFP (green), Par-6::iRFP (magenta) and mCherry-FKBP-PJ/Sac/INPP5E/PJ- dead (cyan) are shown under each quantification figure. For each quantification, means ± standard error of the mean from 20-30 cells pooled across three independent experiments were plotted. **(C)** Purified GST-PKCζ/Par-6::His6 complex, but not purified GST-PKCζ alone or GST-PKCζ^KR8Q^/Par-6::His6 complex, bound to PI4P- and PIP2-liposomes. Scale bars: 5µm (B, all panels)

**Figure 5.**
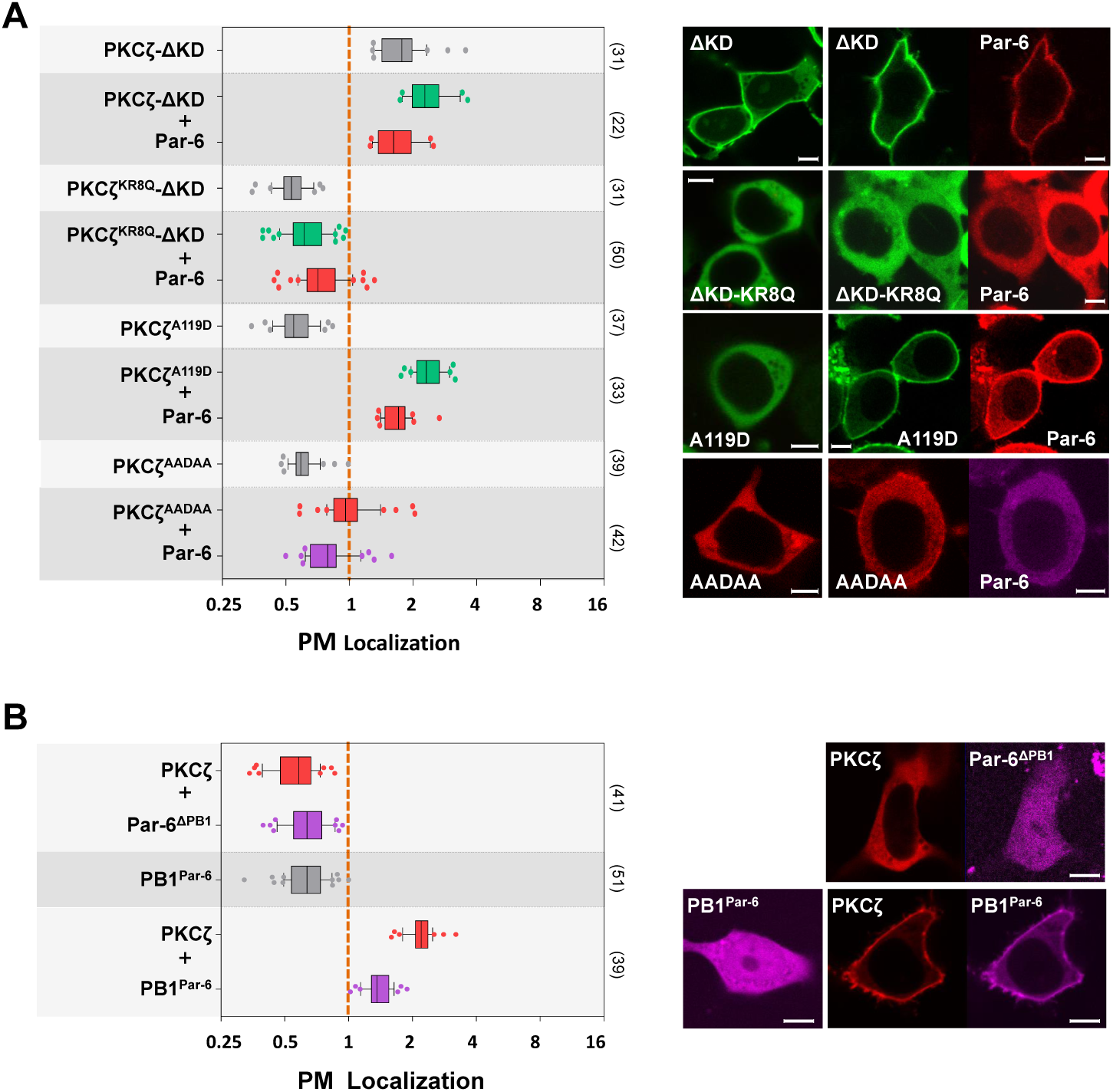
Par-6 interaction with PKCζ is required for polybasic PSr to bind PM. **(A)** PKCζ-ΔKD::GFP (“ΔKD”) localized to PM with our without the co-expression of Par-6::RFP. PKCζ-ΔKD^KR8Q^::GFP (“ΔKD-KR8Q”) was cytosolic with or without the co-expression of Par- 6::RFP. PKCζ^A119D^::GFP (“A119D”) and PKCζ^AADAA^::GFP (“AADAA”) were cytosolic when expressed alone. When co-expressing with Par-6::RFP, PKCζ^A119D^::GFP was strongly PM localized whereas PKCζ^AADAA^::GFP was barely PM localized. **(B)** PKCζ::RFP and Par-6^ΔPB1^::iRFP remained in cytosol when co-expressed. PB1^Par-6^::iRFP was cytosolic when expressed alone, but when co-expressed with PKCζ::RFP both became PM-localized. All experiments were carried out in HEK293 cells. Scale bars: 5µm.

Our data suggest that PI4P and PIP2 likely act redundantly to bind PKCζ/Par-6 complex to PM. This is in contrast to Lgl which appears to rely more on PIP2 for its PM targeting (Dong et al., 2015). Consistently, when we used the similar rapamycin-inducible system to acutely deplete PIP2 in *Drosophila* follicular epithelial cells, Par-6::GFP remained on the PM (Fig. S3) whereas Lgl::GFP showed significant loss from PM in PIP2-depleted cells (Dong et al., 2015). We could not carry out assays by inducible depletion of PI4P or both PI4P and PIP2 as corresponding genetic tools are currently unavailable in *Drosophila*.

Finally, to further confirm the direct binding between aPKC/Par-6 complex and membrane PI4P and PIP2, we carried out liposome binding assays using purified PKCζ, PKCζ/Par-6 complex or PKCζ^KR8Q^/Par-6 complex. As shown in Fig. 4C, purified PKCζ protein alone did not bind PI4P- or PIP2-liposomes, while purified PKCζ/Par-6 complex co-sedimented with PI4P/PIP2-liposomes. Furthermore, neither PKCζ^KR8Q^ nor Par-6 showed binding to PI4P/PIP2-liposomes when purified PKCζ^KR8Q^/Par-6 was used (Fig. 4C). Such in vitro data demonstrate that only aPKC/Par-6 complex, but not aPKC alone, can directly bind membrane PI4P and PIP2 in a polybasic PSr-dependent manner.

### PM-binding of PSr in aPKC is likely allosterically regulated by Par-6

Why is PKCζ alone incapable of binding to PM? Previous studies suggested that the PSr in PKCζ binds the kinase domain (KD) to autoinhibit its kinase activity, and that binding of Par-6 likely induces an allosteric conformation change in PKCζ displacing the PSr from the kinase domain (Graybill et al., 2012). We therefore postulate that in unbound aPKC its PSr is occluded by the kinase domain from binding to the PM but is allosterically exposed once Par-6 binds aPKC. To test this hypothesis, we generated two KD-deletion mutants PKCζ-ΔKD and PKCζ^KR8Q^-ΔKD. In contrast to full length PKCζ, PKCζ-ΔKD localizes to PM in the absence of Par-6 but its PM localization remains dependent on the positive charges of PSr region, as non-polybasic PKC^KR8Q^-ΔKD is cytosolic regardless the presence of Par-6 (Fig. 5A). In addition, merely reducing the interaction between PSr and kinase domain is not sufficient to make PSr accessible to PM binding, as two mutants, PKCζ^A119D^ and PKCζ^AADAA^ carrying previously characterized mutations (Fig. 1A) shown to reduce the autoinhibition of PSr to the kinase domain (Graybill et al., 2012), were all cytosolic (Fig. 5A). When co-expressed with Par-6, PKCζ^A119D^ strongly localized to PM (Fig. 5A), whereas PKCζ^AADAA^ showed barely detectable PM localization likely due to significantly reduced positive charges of PSr by four Arg->Ala mutations (Fig. 1A). Finally, expressing Par-6 PB1 domain (“PB1^Par-6^”), but not Par-6^ΔPB1^, induced PM localization of both PKCζ and PB1^Par-6^ (Fig. 5B), suggesting that interaction between PKCζ and Par-6 PB1 domain alone is both sufficient and necessary to target PKCζ to PM.

Although the lack of full protein structures of aPKC and Par-6 makes it difficult to conduct comprehensive structure-based experiments to further confirm the allosteric regulation of PSr by Par-6, our data strongly support a model that PB1/PB1 interaction between aPKC and Par-6 is both sufficient and necessary to allosterically displace the polybasic PSr from kinase domain in aPKC, exposing PSr to electrostatic binding to the PM.

### Par-6-dependent PM targeting inhibits PKCζ kinase activity

Binding of Par-6 is considered an essential step in regulating the kinase activity of aPKC, although whether Par-6 activates or inhibits aPKC remains unsettled and may well depend on additional regulators presented in different cell types (Hong, 2018). To investigate how Par-6 regulates PKCζ activity *in vivo*, we established aPKC kinase activity assays in HEK293 cells based on the loss of Lgl PM localization that serves as a sensitive, robust and quantifiable readout for measuring aPKC-phosphorylation of Lgl in live cells. When expressed alone in HEK293 cells mammalian Lgl::GFP showed consistently strong and robust PM localization (Fig. 6A) which was strongly reduced in cells expressing PKCζ but not kinase-dead PKCζ^K281W^ (Fig. 6A). An anti-phospho-Lgl antibody confirmed the phosphorylation of Lgl in HEK293 cells co-expressing PKCζ (Fig. 6C), suggesting that overexpressed PKCζ alone contains basal kinase activity sufficient to phosphorylate Lgl, consistent with the fact that in vitro purified aPKC has ∼10% of activated kinase activity (Zhang et al., 2014). To further confirm that the loss of Lgl PM localization is due to the phosphorylation by PKCζ, we generated a non-phosphorylatable Lgl^S6A^::GFP in which all six conserved phospho-serines were mutated to alanine. As expected, Lgl^S6A^::GFP remained on the PM when co-expressed with PKCζ (Fig. 6A).

We then tested the kinase activity of PM targeted aPKC/Par-6 complex. Strikingly, in HEK293 cells co-expressing Lgl::GFP, PKCζ::RFP and Par-6::iRFP, all three proteins were strongly PM localized (Fig. 6A) and anti-phospho-Lgl antibody failed to detect phosphorylation on Lgl (Fig. 6C). Similar results also seen in cells expressing Lgl::GFP and PKCζ::RFP^-2A-^Par-6::iRFP (Fig. 6A). Thus, binding of Par-6 not only targets PKCζ to PM, but also appears to strongly inhibit its kinase activity. This apparent inhibition of PKCζ kinase activity by Par-6 is not due to the PM localization of PKCζ alone, as we made a Lyn11-PKCζ which contains the constitutive PM-binding domain Lyn11 and found that Lyn11-PKCζ also strongly delocalized Lgl::GFP to the cytosol and was efficiently inhibited by Par-6 (Fig. 6B,C). We also compared the Par-6 inhibition on cytosolic PKCζ^KR8Q^ and PM-bound Lyn11-PKCζ^KR8Q^. Both kinases strongly reduced the PM localization of Lgl and were similarly inhibited by Par-6 (Fig. 6B,C), suggesting that PM localization of PKCζ is not required for the kinase inhibition by Par-6.

Curiously, in polarized MDCK cells overexpressed PKCζ failed to dislocalize Lgl::GFP from PM (Fig. S4B), suggesting that Lgl phosphorylation by aPKC is tightly controlled by additional mechanisms. Nonetheless, our results in non-polarized HEK293 cells suggest that binding of Par-6 not only target aPKC to the PM but also inhibits its kinase activity.

### Crb activates aPKC/Par-6 kinase activity to phosphorylate Lgl

The PB1 domain of Par-6 alone (PB1^Par-6^) was capable of targeting PKCζ to PM (Fig. 5B) but it did not inhibit the phosphorylation of Lgl::GFP by PKCζ (Fig. 7A), nor did Par-6ΔPDZ in which the C-terminal PDZ domain was deleted (Fig. 7A). Our results suggest that the PDZ domains or the C-teriminus of Par-6 could be specifically required for inhibiting PKCζ kinase activity on Lgl (Fig. 7A), which is consistent with the finding that overexpression of Par-6 C-terminus inhibits PKCɩ/λ activity in MDCK cells (Kim et al., 2007). The C-terminus of Par-6 interacts with multiple proteins, including activated Cdc42 which binds the Par-6 CRIB domain and moderately activates kinase activity of aPKC/Par-6 complex in vitro (Yamanaka et al., 2001). However, PM localization of Lgl::GFP remained high in HEK293 cells expressing Cdc42^CA^, PKCζ and Par-6 (Fig. 7B), suggesting that Cdc42^CA^ is not sufficient to activate aPKC in our cell-based assays. Our results are consistent with genetic evidences that *Drosophila* Cdc42 is not required for aPKC to phosphorylate Lgl (Hutterer et al., 2004) and Baz (Walther and Pichaud, 2010) in vivo.

**Figure 6.**
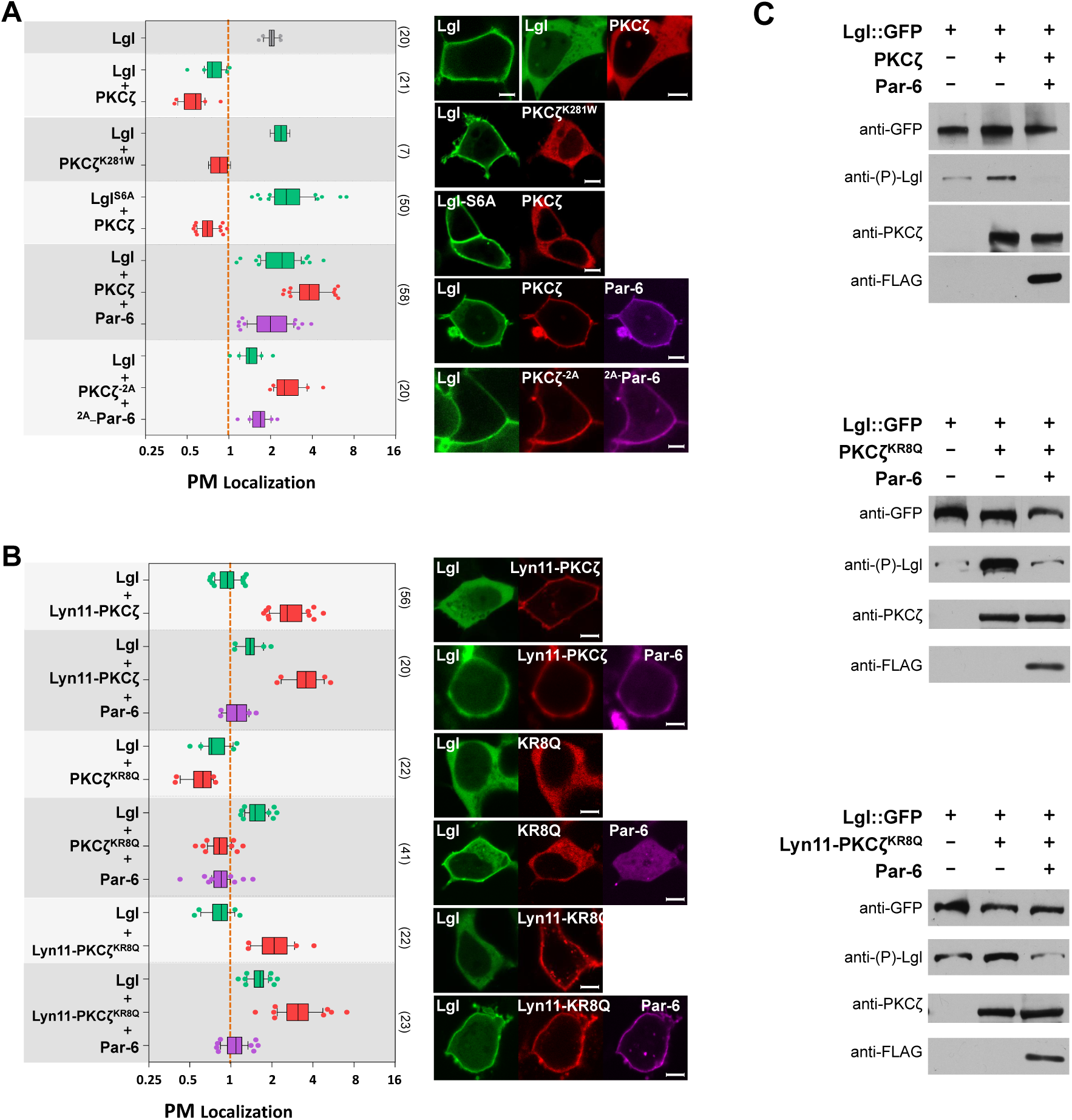
PM-targeted PKCζ/Par-6 complex is inhibited from phosphorylating Lgl. **(A)** PM localization of Lgl::GFP was strongly reduced in cells expressing PKCζ::RFP but not kinase dead PKCζ^K281W^::RFP. Non-phosphorylatable Lgl^S6A^::GFP remained PM localized in cells expressing PKCζ::RFP. Lgl::GFP remained PM localized in cells co-expressing PKCζ::RFP and Par-6::iRFP. Lgl::GFP also showed strong PM localized in cells expressing PKCζ::RFP^-2A-^Par-6::iRFP. **(B)** PM localization of Lgl::GFP was strongly reduced in cells expressing Lyn11-PKCζ::RFP, PKCζ^KR8Q^::RFP (“KR8Q”) and Lyn11-PKCζ^KR8Q^::RFP (“Lyn11-KR8Q”), respectively. In all three cases, co-expression of Par-6::iRFP increased PM localization of Lgl::GFP. **(C)** Cells expressing Lgl::GFP only, expressing both Lgl::GFP and PKCζ::RFP (or PKCζ^KR8Q^::RFP or Lyn11-PKCζ^KR8Q^::RFP), or expressing Lgl::GFP together with PKCζ::RFP (or PKCζ^KR8Q^::RFP or Lyn11-PKCζ^KR8Q^::RFP) and FLAG::Par-6, were directly lysed in SDS-loading buffer and analyzed by western blot. Anti-(P)-Lgl: antibody against phosphorylated Lgl. All experiments were carried out in HEK293 cells. Scale bars: 5µm.

**Figure 7.**
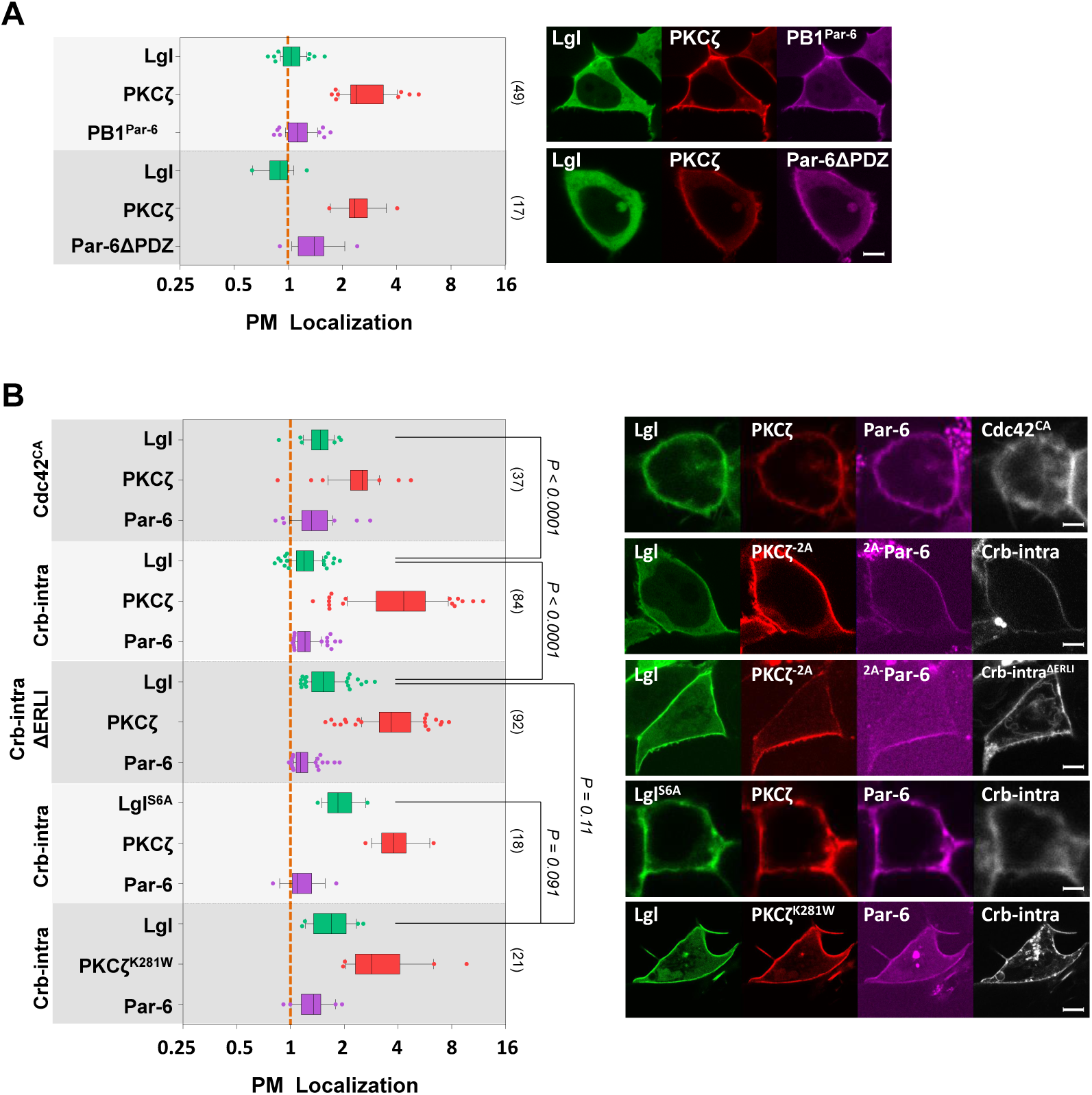
Crb-intra activates PM-targeted aPKC/Par-6 complex. **(A)** In HEK293 cells expressing either PB1^Par-6^::iRFP or Par-6ΔPDZ::iRFP, PKCζ::RFP was strongly PM-localized whereas Lgl::GFP was strongly reduced from PM. **(B)** In HEK293 cells expressing Lgl::GFP, PKCζ::RFP and Par-6::iRFP, PM localization of Lgl::GFP was strongly reduced when BFP::Crb-intra, but not BFP::Crb-intra^ΔERLI^ or BFP::Cdc42^CA^, was co-expressed. Lgl^S6A^::GFP remained on PM in cells expressing PKCζ::RFP, Par-6::iRFP and BFP::Crb-intra. Lgl::GFP remained on PM in cells expressing BFP::Crb-intra, Par-6::iRFP and kinase-dead PKCζ^K281W^::RFP. Scale bars: 5µm.

Besides Cdc42, apical polarity protein Crb also interacts with Par-6. Crb is a transmembrane protein and the C-terminus of its intracellular domain is a PDZ-binding motif (PBD) that can bind the PDZ domain in Par-6 (Kempkens et al., 2006). We therefore investigated whether Crb could directly activate the kinase activity of aPKC/Par-6 complex through its interaction with the Par-6 PDZ domain. Par-6 became localized to the PM in HEK293 cells expressing membrane-bound Crb intracellular domain (Crb-intra) but not Crb-intra^ΔERLI^ in which the C-terminal PBD was deleted, confirming that Par-6 binds to the PBD of Crb-intra (Fig. S4A). Moreover, in HEK293 cells expressing aPKC and Par-6 together with Crb-intra but not Crb-intra^ΔERLI^, Lgl::GFP was strongly reduced from the PM (Fig. 7B). The loss of Lgl::GFP from PM in cells expressing Crb-intra, aPKC and Par-6 is phosphorylation-dependent, as non-phosphorylatable Lgl^S6A^::GFP remained on PM in these cells (Fig. 7B). Lgl::GFP also maintained PM localization in cells expressing Crb-intra, Par-6 and kinase-dead aPKC^K281W^ (Fig. 7B). These results support a model that direct interaction between Crb-intra and Par-6 in a PM-bound aPKC/Par-6 complex activates aPKC kinase activity.

Consistent with our cell-based assays, overexpression of Crb in *Drosophila* follicular cells and embryonic epithelial cells expanded Crb and DaPKC into basolateral PM, and strongly delocalized Lgl::GFP, but not non-phosphorylatable Lgl^S5A^::GFP, from basolateral PM (Fig. 8A-C). The loss of Lgl::GFP from PM in Crb-overexpressing cells was completely reversed when DaPKC or Par-6 was also knocked down by RNAi (Fig. 8E,F). Furthermore, the loss of PM Lgl::GFP in Crb-overexpressing cells is comparable to cells overexpressing DaPKC-ΔN which is considered fully activated due to the deletion of the N-terminus including PSr (Betschinger et al., 2003) (Fig. 8G). Thus, in Crb-overexpressing follicular cells DaPKC is the key kinase phosphorylating Lgl and is likely highly activated. In contrast, in *Drosophila crb^-/-^* embryos DaPKC also extended to the basolateral PM but Lgl remained on PM (Fig. 8D), suggesting that Crb is necessary to promote DaPKC-phosphorylation on Lgl in embryonic epithelial cells in vivo.

**Figure 8.**
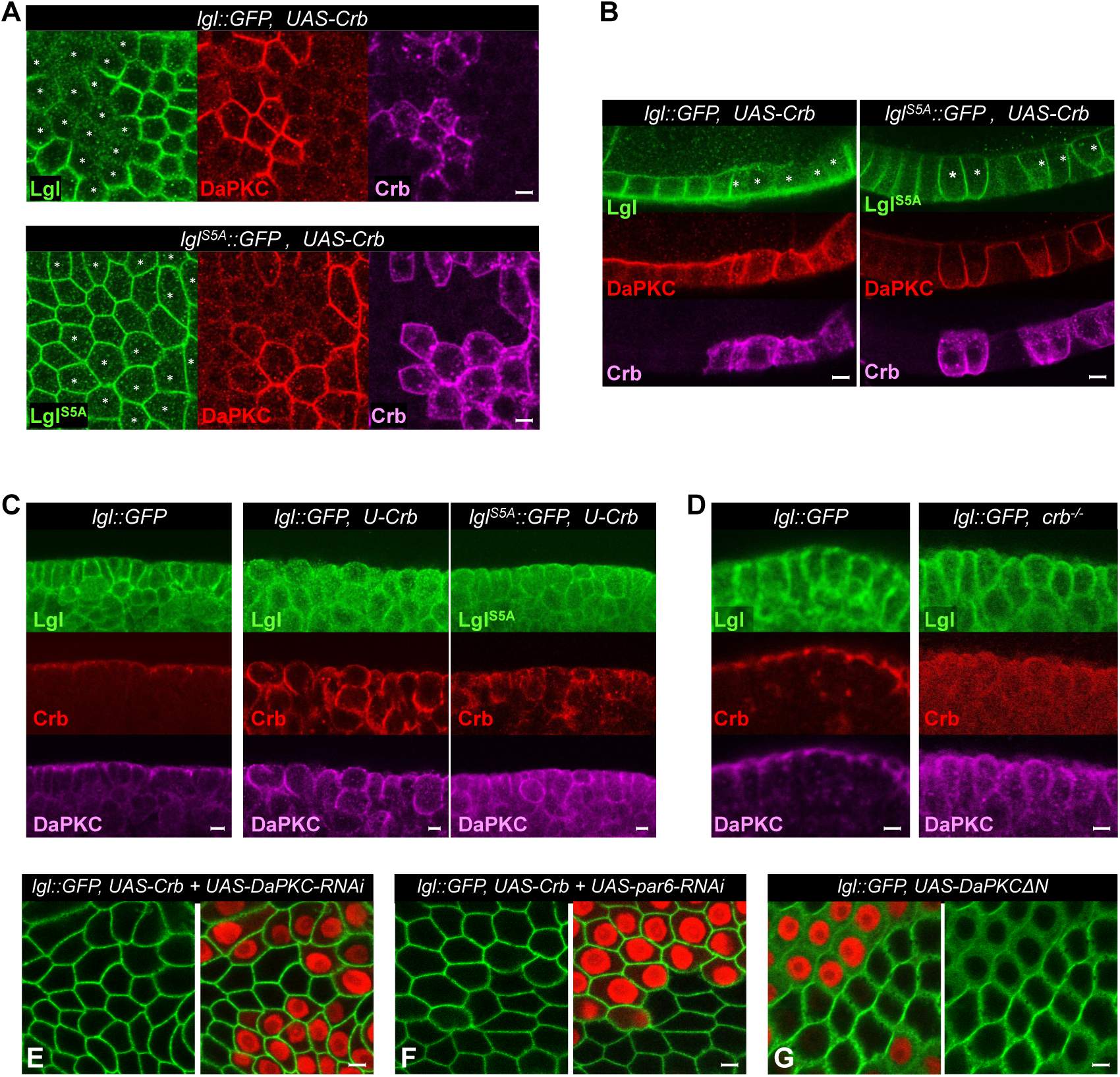
Crb promotes DaPKC phosphorylation on Lgl in vivo. (**A, B**) *Drosophila lgl::GFP* or *lgl*^*S6A*^::*GFP* follicular epithelial cells overexpressing Crb were immunostained for GFP (green), DaPKC (red) and Crb (magenta). Images in A are in tangential view and were sectioned below the apical surface of follicular cells where Crb and aPKC normally are absent. Images in B are in cross-section view of follicular epithelial cells, showing overexpressed Crb expanded into the lateral PM along with DaPKC. Cells overexpressing Crb are highlighted by asterisks in green channel images. **(C)** Wild type *lgl::GFP* embryos and embryos of *lgl::GFP UAS-Crb/Mat-Gal4* or *lgl*^*S5A*^::*GFP UAS-Crb/Mat-Gal4* were immunostained for GFP (green), Crb (red) and aPKC (magenta). All embryonic epithelial cells were in cross-section view. Note the loss of Lgl::GFP, but not Lgl^S5A^::GFP, from the PM under Crb over-expression driven by *Mat-Gal4* (see Materials and Methods). **(D)** Wild type *lgl::GFP* embryos and *lgl::GFP; crb^-/-^* mutant embryos were immnuostained for GFP (green), Crb (red) and aPKC (magenta). In *crb*^-/-^ embryos red channel was overexposed to confirm no detectable expression of Crb. In *crb*^-/-^ embryonic epithelial cells both Lgl and DaPKC became localized all around PM. (**E, F**) Lgl::GFP remained on PM in Crb-overexpressing cells that were also expressing *DaPKC-RNAi* (E) or *par-6-RNAi* (F). (**G**) Lgl::GFP was severely lost from PM in follicular cells expressing DaPKCΔN. In E-G: cells expressing *DaPKC-RNAi, par-6-RNAi* or DaPKCΔN are marked by RFP expression (see Materials and Methods). Scale bars: 5µm.

## DISCUSSION

### Electrostatic binding to phosphoinositides by polybasic PSr targets aPKC to PM

In this report we show that the pseudosubstrate region (PSr) in aPKC is a typical polybasic domain capable of directly targeting aPKC to PM via its electrostatic interaction with negatively charged phosphoinositides PI4P and PIP2 in PM. This is in contrast to the assumption that protein-protein interactions are solely responsible for localizing aPKC to PM or cell cortex. In addition, unlike phosphorylatable polybasic domains in Lgl, Numb and Miranda, the polybasic PSr in aPKC has not been shown phosphorylatable. Instead we report a novel example of a potential allosteric regulation of a polybasic domain for PM-binding. As illustrated in Figure 9, our data support a model that the polybasic PSr in unbound aPKC is occluded by the kinase domain from binding PM, whereas the binding of Par-6 to aPKC via PB1 domains induces potential conformational changes in aPKC that make the polybasic PSr accessible to PM-binding.

**Figure 9.**
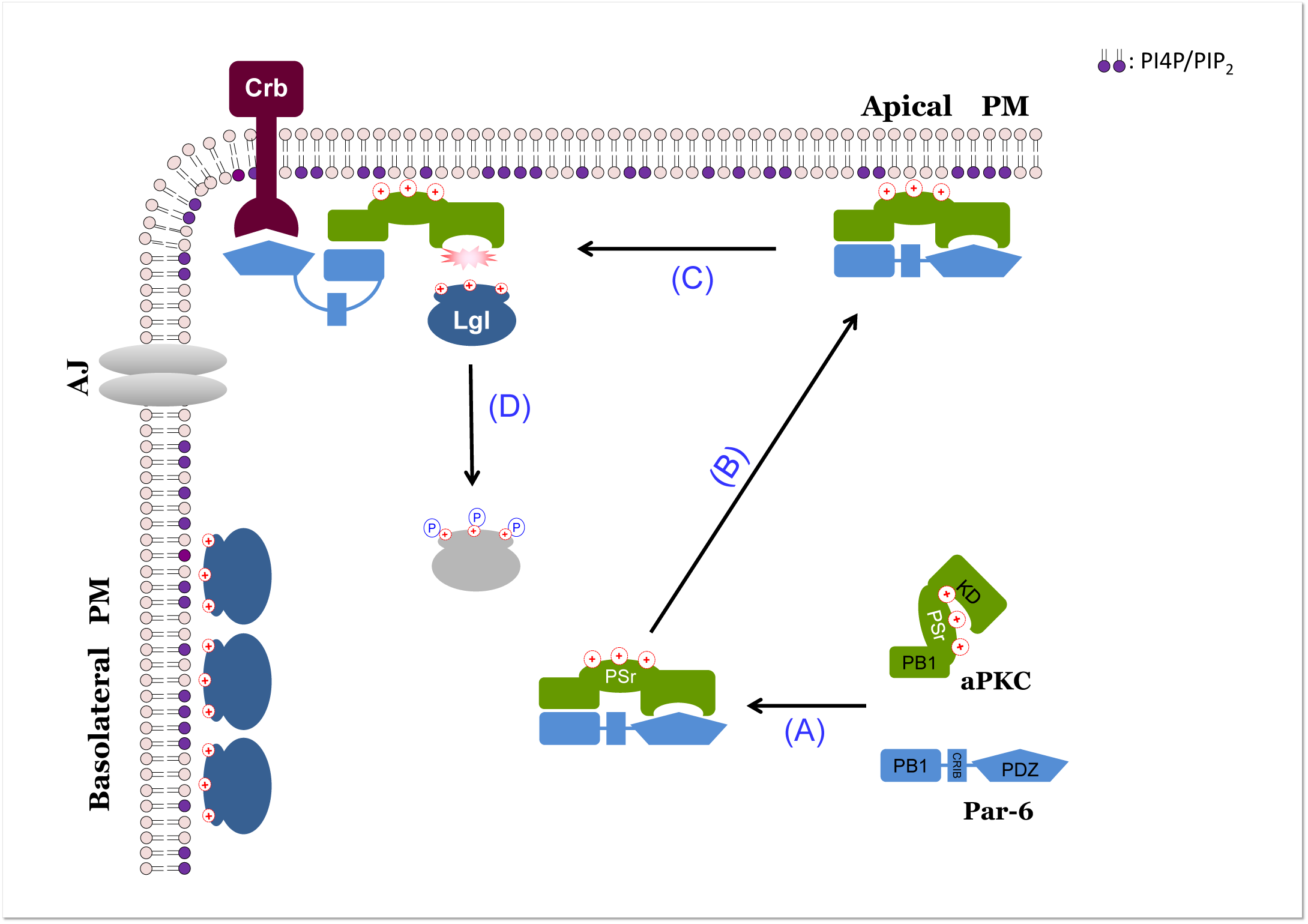
A hypothetic model of aPKC PM targeting and kinase activation. **(A)** Free cytosolic aPKC in auto-inhibited conformation has polybasic PSr blocked by the kinase domain from binding to PM. Binding of Par-6 to aPKC induces conformation changes that expose the PSr in aPKC and allows the C-terminus of Par-6 to simultaneously inhibit the aPKC kinase domain. **(B)** Polybasic PSr in aPKC/Par-6 complex binds to PM via electrostatic interaction with PI4P and PIP2 which are uniquely enriched on PM. **(C)** Intracellular domain of apical polarity protein Crb interacts with the C-terminal PDZ domain of Par-6 and releases its inhibition on aPKC kinase domain. Interaction with Crb could also facilitate the apical-enrichment of PM-bound aPKC/Par-6 in cells. **(D)** Activated aPKC phosphorylates Lgl to prevent it from binding to apical PM. Illustration is based on *Drosophila* epithelial cells. AJ: adherens junction. PM: plasma membrane. PSr: pseudo-substrate region.

Cdc42 plays an important role in mediating the PM targeting of aPKC/Par-6 in certain cell types such as *C. elegans* one-cell embryos, and we found that activated Cdc42 binds Par-6 which in turn can recruit non-polybasic aPKC^KR8Q^ to the PM in HEK293 cells. However, our in vitro and in vivo studies suggest that electrostatic binding to PI4P and PIP2 alone can be sufficient to localize aPKC/Par-6 complex to the PM. Our results are consistent with findings that removing Cdc42 in *C. elegans* late embryos or in *Drosophila* pupal epithelial cells does not severely disrupt aPKC and Par-6 localization (Georgiou et al., 2008; Zilberman et al., 2017). Whereas Cdc42 directly regulates aPKC/Par-6 PM targeting under certain cellular contexts, our results demonstrate a PM targeting mechanism of aPKC/Par-6 that is mechanistically independent of Cdc42 or other aPKC/Par-6 interacting proteins.

Our hypoxia assay suggest that at least in *Drosophila* follicular and embryonic epithelial cells, binding to PI4P and PIP2 is likely the primary mechanism localizing aPKC/Par-6 to the PM in vivo. In these cells the Par-6-dependent electrostatic binding to PM by PSr likely functions as the first step to localize aPKC/Par-6 to the PM, followed by further enrichment to specific membrane domains such as apical PM through protein-protein interactions with, for instance, apical Crb complex. Thus, while mechanisms governing the aPKC/Par-6 subcellular localization can be heavily cell-type specific and multi-layered, our results highlight that the electrostatic PM-binding property of aPKC needs to be taken into consideration when studying the protein regulators of aPKC/Par-6 subcellular localization.

Our studies also suggest potential new negative regulators that may act specifically at the electrostatic interaction to prevent aPKC from binding to the PM, by mechanisms such as masking the PSr, inhibiting the allosteric changes induced by Par-6 or sequestering the phospholipids in the PM. Moreover, it is notable that a PI4P- and PIP2-dependent mechanism makes aPKC PM targeting vulnerable to stress conditions such as hypoxia and ATP inhibition that deplete these phospholipids in the PM (Dong et al., 2015). The acute and reversible loss of PM targeting of polybasic polarity proteins like Lgl, aPKC and Par-6 may have profound implications in epithelial cells for maintaining and restoring their apical-basal polarity when undergoing hypoxia, ischemia and reperfusion.

### Par-6 controls both aPKC PM targeting and kinase activity

Mechanisms coupling the aPKC kinase activity with its subcellular localization are essential for aPKC to phosphorylate targets at the right place and right time (Hong, 2018), but molecular details about these mechanism remain largely elusive. Recent studies began to reveal exciting details on how aPKC/Par-6 kinase activity and subcellular localization can be regulated by the clustering of Par-3(Baz) and diffusive interactions with Cdc42 (Dickinson et al., 2017; Rodriguez et al., 2017), but these studies so far have been limited to the process of anterior-posterior (A-P) polarization in worm one-cell embryos. Here we show that, in *Drosophila* epithelial cells and cultured mammalian cells, the electrostatic binding of aPKC to PM alone may provide a mechanism for Par-6 to play a pivotal role in coupling the PM targeting and control of kinase activity of aPKC (Fig.9). It is possible that conformational changes in aPKC induced by Par-6 may not only expose PSr to PM-binding but also allow the C-terminus of Par-6 to simultaneously inhibit aPKC’s kinase domain. More experiments, however, are needed to further validate this hypothesis.

Our allosteric model is based on the previous studies suggesting the displacement of PSr from aPKC upon binding of Par-6 (Graybill et al., 2012). However, in that study Par-6 appears to activate aPKC in vitro and in *Drosophila* S2 cells. Such discrepancy could be due to the fact that S2 cells already express proteins capable of releasing the inhibition of Par-6 on aPKC. It should be noted that studies from multiple groups yielded conflicting results on whether Par-6 inhibits or activates aPKC kinase activity (Atwood et al., 2007; Chabu and Doe, 2008; Graybill et al., 2012; Lin et al., 2000; Yamanaka et al., 2001). Most of these studies relied on immunoprecipitated or reconstituted aPKC/Par-6 complex to measure the kinase activity in solution in vitro. Given that the majority of aPKC substrates are PM-bound polybasic domain-containing proteins, it might be critical to assay aPKC/Par-6 kinase activity in its PM-bound form in live cells.

In addition, it is conceivable that binding to PM may physically shield the polybasic domain in a target protein from being accessible to aPKC, as suggested by the increased resistance of membrane-bound Lgl to aPKC-phosphorylation in vitro (Visco et al., 2016). This is consistent with our finding that phosphorylation of Lgl appear to be inhibited when both Lgl and aPKC/Par-6 complex are electrostatically attached to PM. On the other hand, the transient and dynamic nature of lipid-binding by polybasic domains (Hammond et al., 2009) could also effectively enrich a local cytosolic pool of target proteins near PM, which works in favor of PM-bound aPKC to encounter its substrates. Such intricate relationships between PM-targeting and aPKC-phosphorylation of polybasic polarity proteins remain to be further explored.

### PM targeting and spatial control of aPKC kinase activation

Similar to cPKC and nPKC, the electrostatic binding of PSr to PM could potentiate the kinase activity of aPKC/Par-6 complex by further preventing PSr from autoinhibiting the kinase domain (Garg et al., 2013; Rosse et al., 2010). This is supported by the in vitro studies that Par-6 binding to aPKC inhibits its kinase activity but also potentiates its activation upon subsequent molecular events (Yamanaka et al., 2001). PM-bound aPKC/Par-6 complexes with inhibited but potentiated kinase activity are ideal targets for additional activation to fine-tune aPKC localization and kinase activity. Curiously, we found that Cdc42 did not activate PM-bound aPKC/Par-6 to phosphorylate Lgl in our cultured cell assays, but interaction between Par-6 and apical transmembrane protein Crb reversed the Par-6’s inhibition on aPKC kinase activity. Supporting the role of Crb in enriching and activating aPKC/Par-6, Crb colocalizes with aPKC/Par-6 complex and is required for enriching the aPKC/Par-6 complex to the apical PM in fly embryonic epithelial cells and in many specialized membranes in *Drosophila* including apical membranes of photoreceptors (Hong et al., 2003; Pellikka et al., 2002) and lumens in scolopale cell in chondotoral organs (Y.H., data not shown). However, Crb is not required for apical enrichment of aPKC in follicular cells (Sherrard and Fehon, 2015), therefore additional proteins besides Crb must be responsible for the apical enrichment and activation of aPKC/Par-6 in different types of epithelial cells.

It is of note that our studies are primarily based on cultured cells and limited cell types of *Drosophila* epithelial cells, thus present a simplified model regarding the relationships between aPKC/Par-6 and its regulators. Our data do highlight that aPKC/Par-6 activity can be regulated by multiple regulators such as PM phosphoinositides, Cdc42 and Crb under different cellular and polarity contexts. Extrapolating our study to further investigate the role of aPKC in regulating the cell polarity requires experiments to take into account the more complicated cellular contexts in different cell types.

## ACKNOWLEDGEMENTS

We are grateful to Drs. Jane Wang, Gerald Apodoca, Mark Peifer, Eli Knust, Thomas Weide, David Bilder, Tony Harris, Daniel St Johnston, Jean Wilson, and Ricardo Biondi for reagents and fly stocks, Lu Jiang and Kriti Sanghi for technical assistances, Dr. Marijn Ford for helps on protein purifications, Dr. Simon Watkins and University of Pittsburgh Medical School Center for Biologic Imaging for generous imaging and microscopy support, Bloomington Stock Center for fly stocks, and Developmental Studies Hybridoma Bank (DSHB) for antibodies.

## COMPETING INTERESTS

The authors declare no competing or financial interests.

## AUTHOR CONTRIBUTIONS

Conceptualization: Y.H., W.D., G.R.H.; Investigation: W.D., J.L., X.Z., Y.W., K.L., G.R.H. and Y.H.; Writing - Review &Editing: Y.H., W.D., G.R.H.; Funding acquisition: Y.H., G.R.H.; Supervision: Y.H.

## FUNDING

This work was supported by grants NIH-NCRR R21RR024869 (Y.H.), NIH-NIGMS R01GM086423 and R01GM121534 (Y.H.), NIH 1R35GM119412-01 (G.R.H.). University of Pittsburgh Medical School Center for Biologic Imaging is supported by grant 1S10OD019973-01 from NIH.

## MATERIALS AND METHODS

### Fly stocks and genetics

Following *Drosophila* stocks were used this study: knock-out alleles of *lgl*^*GX*^ (“*lgl*^*KO*^”) and *crb*^*GX*^ (“*crb*^*KO*^”) and knock-in alleles of *lgl::GFP* and *lgl*^*S5A*^::*GFP* were previously described (Dong et al., 2015; Huang et al., 2009). *w; UAS-Crb*; (gift from Dr. Eli Knust, Dresdon, Germany), *w par-6^Δ226^ FRT^9-2^/FM6* (a gift from Dr. Jurgern Knoblich, IMP, Vienna, Austria), *y w; DaPKC^k06403^/CyO* (BL#10622), *w; His2Av::mRFP* (BL#23651), *y w ubi-GFP^NLS^ FRT^9-2^* (BL#5154), *w; Act5C(FRT.CD2)-Gal4, UAS-RFP/TM3, Sb* (BL#30558), *w; FRT^G13^ DaPKC^k06403^/CyO; ubi-DaPKC::GFP/TM6, w; FRT^G13^ DaPKC^k06403^/CyO; ubi-DaPKC^KR8Q^::GFP/TM6, w par-6::GFP hs-FLP par-6^Δ226^ FRT^9-2^/FM7C, w; lgl::GFP hs-FLP^38^; Act5C(FRT.CD2)-Gal4, UAS-RFP/TM3, Sb, w; lgl::GFP UAS-Crb/CyO, w, lgl^S5A^::GFP UAS-Crb/CyO, w; lgl^S5A^::GFP/CyO; hs-FLP Act5C(FRT.CD2)-Gal4, UAS-RFP/TM3, w hs-FLP; FRT^G13^ His2Av::mRFP, w par-6::GFP hs-FLP par-6^Δ226^ FRT^9-2^/FM7C; lgl^GX^ FRT^40A^/CyO, w par-6::GFP hs-FLP par-6^Δ226^ FRT^9-2^/FM7C; ubi-RFP^NLS^ FRT^40A^/CyO, w; lgl::GFP UAS-aPKC-CAAX (UAS-aPKC-CAAX* is a gift from Dr. Sonsoles Campuzano, Univerisidad Autonoma de Madrid, Spain), *w UASp>mRFP::FKBP-5Ptase* (“FKBP-INPP5E”) and *w;; UASp>Lck-FRB::CFP* are gifts from Dr. De Renzis, EMBL Heidelberg, Germany (Reversi et al., 2014), *w; αTub67C-Gal4^V2H^; αTub67C-Gal4^V37^* (“Mat-Gal4”, a gift from Dr. Mark Peifer, University of North Carolina at Chapel Hill, USA) (Schaefer et al., 2018), *w; GFP::aPKC/CyO* (gift from Dr. Daniel St Johnston), *w; UAS-Cdc42^CA^* (BL#4854), *UAS-cdc42-RNAi, w;* (BL#29004), *y, v; UAS-DaPKC-RNAi* (BL#25946), *y, v; UAS-Par-6-RNAi* (BL#39010), *y, v; UAS-crb-RNAi* (BL#38373), *w;; UAS-DaPKC-CAAX, w;; UAS-DaPKC-CAAX, UAS-Par-6; w;; UAS-Par-6* (gifts from Dr. Tony Harris).

Transgenic flies of *ubi-DaPKC::GFP, ubi-aPKC^KR8Q^::GFP* and *par-6::GFP* were generated by phiC31-mediated integration protocol (Huang et al., 2009). *attP*^*VK00033*^ (BL#24871) stock was used to integrate ubi-DaPKC::GFP and ubi-DaPKC^KR8Q^::GFP constructs into the 3^rd^ chromosome, while *attP*^*ZH2A*^ (BL#24480) stock was used to integrate par-6::GFP constructs into the X chromosome. *par-6::GFP* was further recombined with *par-6^Δ226^* null allele to generate *w par-6::GFP par-6^Δ226^* of which homozygote females and hemizygote males are fully viable and fertile, indicating a complete rescue of *par-6^Δ226^* by *par-6::GFP*.

*Drosophila* cultures and genetic crosses are carried out at 25°C. Detail information about the strains from Bloomington stock center can be found in FlyBase.

### Molecular cloning

To make ubi-aPKC::GFP, ubiquitin promoter (1872bp) was PCR from plasmid pWUM6 (a gift from Dr. Jeff Sekelsky, University of North Carolina at Chapel Hill) using primers 5-AGTGTC GAATTC CGCGCAGATC GCCGATGGGC and 5-CTGGAC GCGGCCGC GGTGGATTATTCTGCGGG and inserted into pGE-attB vector (Huang et al., 2009) to generate vector pGU. DNA fragments encoding aPKC::GFP and aPKC^KR8Q^::GFP were then inserted into pGU vector. To make *par-6::GFP*, a 4.3kb *par-6* genome DNA including 1Kbp upstream and 250bp downstream sequences was PCR-amplified from *Drosophila* genomic DNA using primers 5-ATGCGGCCGC GCTCTTCGGC TCTCGGATAG TCG and 5-GACGCGTGAT TAAGGCCCGG CTAATG, subcloned into pGE-attB vector. AvrII enzyme site was added before stop code for GFP insertion. More details about DNA constructs used in this report are listed in Supplementary Table S1. NCBI RefSeq ID: Par-6 (human): NP_058644.1, Lgl (mouse): NP_001152877.1. Plasmids containing PKCζ and PKCι coding sequences were gifts from Dr. Jane Wang (U of Pittsburgh)

### Live imaging and hypoxia treatment of aPKC::GFP and Par-6::GFP in *Drosophila* -epithelial cells

Embryos and dissected ovaries were imaged according to previously described protocol (Dong et al., 2015; Huang et al., 2011). Embryos were collected over night at 25°C. Ovaries from adult females of several days old were dissected in halocarbon oil (#95). Dechorinated embryos or dissected ovaries were mounted in halocarbon oil on an air-permeable membrane (YSI Membrane Model #5793, YSI Inc, Yellow Springs, OH) that is sealed by vacuum grease on a custom-made plastic slide over a 10mmx10mm cut-through window. After placing the coverslip on top, membrane at the bottom ensures the sufficient air exchange to samples during the imaging session. The slide was then mounted in an air-tight micro chamber (custom made) for live imaging under confocal microscope. Oxygen levels inside the chamber were controlled by flow of either air or custom O_2_/N_2_ gas at the rate of approximately 1-5 cc/sec. Images were captured at room temperature (25°C) on a Leica TCS-NT confocal microscope (PL APO 40x oil objective, NA=1.25) by Leica TCS-NT software, or an Olympus FV1000 confocal microscope (40x Uplan FL N oil objective, NA=1.3) by Olympus FV10-ASW software, or a Nikon A1 confocal microscope (Plan Fluo 40x oil objective, NA=1.3) by NIS-Elements AR software. Images were further processed in ImageJ and Adobe Photoshop.

### Purification of PKCζ and PKCζ/Par-6 complex from cultured cells

GST-PKCζ (gift from Dr. Ricardo Biondi (Zhang et al., 2014)), GST-PKCζ^KR8Q^, and Par-6::Hisx6 were expressed in Expi293F cells via transfection using the ExpiFectamine Transfection Kit (Gibco). Approximately 22.5×10^7^ suspension-adapted Expi293F cells were diluted in Expi293 Expression Medium to the final volume of 76.5 ml and cultured in a 37°C shaker (Eppendorf New Brunswick S41i). 90µg of plasmid DNA was mixed with 240µl Expifectamine reagent into a total final volume of 9ml OPTI-MEM medium and was incubated for 25 min. The mixture was then added to Expi293F cells to a total volume of 85.5 ml. After 18 hours of incubation in the 37°C shaker, 450µl of Transfection Enhancer I and 4.5 ml of Transfection Enhancer II were added to the cells. Cells were harvested after another 18 hrs of incubation, resuspended in 50ml PBS buffer, and homogenized by ice-cold tight fitting Dounce homogenizer. To purify PKCζ/Par-6 or PKCζ^KR8Q^/Par-6 complex, cells expressing Par-6::His6 were mixed with equal volume of cells expressing GST-PKCζ or GST-PKCζ^KR8Q^ and homogenized. After centrifuging, supernatant from lysate were collected for purification by Pierce Glutathione Agarose. Bacteria BL21 was used to express and purify GST and GST fusion of aPKC PSr proteins as previously described (Dong et al., 2015).

### Liposome pull-down assays

Liposomal binding assays were carried out as described (Kim et al., 2008). To prepare liposomes, lipid mixture of 37.5% PC (Cat#840051C), 10% PS (Cat#840032C), 37.5% PE (Cat#840021C), 10% Cholesterol (Cat#700000P) and 5% PI(4,5)P_2_ (Cat#840046X) or PI4P (Cat#840045X, all lipids were purchased from Avanti Polar Lipids Inc) was dried and resuspended to a final concentration of 1 mg/ml of total phospholipids in HEPES buffer. After 30 min sonication, formed liposomes were harvested at 16,000 × g for 10 min and resuspended in binding buffer (HEPES, 20 mM, 7.4, KCl 120 mM, NaCl 20mM, EGTA 1mM, MgCl 1mM BSA 1mg/ml). In each liposome-binding assay, approximately 0.1 μg of purified protein or protein complex was mixed with 50μl of liposome suspension. After 15 min incubation at room temperature, liposomes were pelleted at 16,000 × g for 10 min and were analyzed by western blot to detect co-sediment of target protein(s).

### Cell culture and imaging

HEK293 cells were cultured in glass bottom dishes (In Vitro Scientific) and were transfected with DNA using X-treme Gene 9 DNA transfection reagent (Sigma Cat# 6365787001). After 24 to 40 hours of transfection cells were mounted and imaged on an Olympus FV1000 confocal microscope (40x Uplan FL N oil objective, NA=1.3) by Olympus FV10-ASW software, or a Nikon A1 confocal microscope (Plan Fluo 40x oil objective, NA=1.3) by NIS-Elements AR software. For images to be used for quantification, parameters were carefully adjusted to ensure no or minimum over-exposure. In addition, when necessary a fluorescent PM dye (CellMask DeepRed Plasma Membrane Stain, ThermoFisher, Cat#C10046) was added the cell culture prior to live imaging to help visualizing the PM for later quantifications. Madin-Darby canine kidney (MDCK) II cells were cultured in MEMα media containing 10% fetal bovine serum (Gibco) and 1% penicillin and streptomycin (Gibco). For growing polarized monolayers, MDCK cells were cultured on 0.4µm Transwell filters (Corning) for 3 days. Transfection assay were carried out on day 4 using Lipofectamine 2000 (Thermo Fisher Scientific) and cell were imaged on day 5.

### Quantification of PM localization

PM localization were measured in Image J by custom macro scripts. For each image, PM masks were generated by an à trous waveleta decomposition method (Hammond et al., 2014; Olivo-Marin, 2002) base on the channel that either contains PM-localized proteins or fluorescent PM dyes. Cytosol masks were generated by segmentation using threshold based on the mean pixel value of the ROI. Cells expressing all transfected fluorescent proteins were selected for measurement by drawing ROIs around each cell. Due to the use of computer generated PM and cytosol masks, the exact shape of the ROI was not critical except that PM segments in contact with neighboring expressing cells were avoided. Custom macros were used to automatically measure PM and cytosolic intensities of each fluorescent protein in each cell marked by ROI in the sample image. Background were auto-detected by the macro based on the minimal pixel value of the whole image. The PM localization index for each fluorescent protein was calculated by the macro as the ratio of [PM - background]/[cytosol - background]. Data were further processed in Excel, visualized and analyzed in Graphpad Prism.

### Biochemistry

For Lgl-phosphorylation assay, HEK293 cells were cultured in DMEM supplemented with 10% FBS. 24 hours after transient transfection, cells were directly lysed in SDS loading buffer and equal volumes of cell lysates were resolved in 12% SDS-PAGE. Proteins were detected by western blot using antibody chicken anti-GFP, 1:5,000 (Aves Lab, Cat# GFP-1020), rabbit anti-phospho-mLgl, 1:1000(Abgent, Cat# AP2198a), rabbit anti-PKC, 1:5000 (“pan-aPKC” antibody, Santa Cruz, Cat# Sc-216), PKCζ-specific antibody (C24E6) rabbit mAb (Cell Signalling, Cat #9368), or mouse anti-Flag 1:5,000 (Sigma, Cat# F3165). For immunoprecipitation experiments, cells were lysed in RIPA buffer (25mM Tris-HCl, pH 7.4, 150 mM NaCl, 0.1% SDS, 0.5% sodium deoxycholate, 1% Triton X-100). Protein G-Sepharose beads were incubated with home-made and affinity-purified rabbit anti-GFP antibody (Huang et al., 2009) for 1 hour followed by incubation with the equal amount of each lysate for 1 hour. The beads were washed and boiled with SDS loading buffer. Supernatants were detected by western blot.

### Induction of FKBP12-phosphatase and Lyn11-FRB::CFP dimerization in HEK293 cells

The procedure has been described in detail previously (Hammond et al., 2014). In brief, HEK293 cells cultured in 35 mm glass bottom dishes (In Vitro Scientific) were transiently transfected with 1 µg total DNA which included the Lyn11-FRB::CFP recruiter, FKBP12-phosphatases (Hammond et al., 2012), PKCζ::GFP and Par-6::iRFP as indicated. After 22-26 hours, cells were imaged in Fluoro-Brite medium (Life Technologies) using a Nikon A1R confocal laser scanning microscope though a 100x, NA/1.45 plan apochromatic objective lens. Time lapse imaging started 2 minutes prior to bath addition of 1 µM rapamycin. CFP::Lyn11-FRB image images were used to generate binary masks to define PM. Plasma membrane localization of each reporter was then calculated from the ratio of fluorescence within the PM to the whole cell, and is expressed relative to the average before rapamycin addition (Hammond et al., 2014).

### Induction of mRFP::FKBP-5Ptase and Lck-FRB::CFP dimmerization in live *Drosophila* follicular cells

Young females of *w UASp>mRFP::FKBP-5Ptase / w par-6::GFP par-6^Δ226^; hs-FLP Act5C(FRT.CD2)-Gal4 UAS-RFP/UASp>Lck-FRB::CFP* or *w UASp>mRFP::FKBP-5Ptase / +; lgl::GFP hs-FLP /+; Act5C(FRT.CD2)-Gal4 UAS-RFP / UASp>Lck-FRB::CFP* were heat-shocked at 37°C for 1hr. Ovaries were dissected 4 days later in 1xPBS prior, mounted in a drop of 20μl Schneider’s medium containing 10μM rapamycin on a gas-permeable slide and imaged live, as previously described (Huang et al., 2011) (Dong et al., 2015). Treated *lgl::GFP* ovaries served as a positive control (Dong et al., 2015).

### Generation of mitotic mutant clones in *Drosophila* follicular epithelia

Mutant follicular cell clones of *lgl*^*KO*^ or *aPKC*^*k06403*^ were generated by the routine FLP/FRT technique. Young females were heat-shocked at 37°C for 1 hour and their ovaries were dissected 3 days later.

### Immunostaining and confocal imaging

Immunostaining of embryos and adult ovaries were carried out as described (Huang et al., 2009). Primary antibodies: rabbit anti-GFP (Huang et al., 2009) 1:1500; chicken anti-GFP (Aves Lab) 1:1000; rabbit anti-Lgl (d-300, Santa Cruz) 1:200; rabbit anti-aPKC (Santa Cruz) 1:1000. Secondary antibodies: Cy2-, Cy3 or Cy5-conjugated goat anti-rabbit IgG, anti-chicken IgG, goat anti-rat IgG, goat anti-mouse IgG and goat anti-guinea pig IgG (The Jackson ImmunoResearch Lab), all at 1:400. Images were collected on an Olympus FV1000 confocal microscope and processed in Adobe Photoshop and ImageJ.

### Genotypes of *Drosophila* Samples in Figures

**Figure 1:** (**C**) *w; FRT^G13^ DaPKC^k06403^ / CyO; ubi-DaPKC::GFP / TM6; w; FRT^G13^ DaPKC^k06403^ / CyO; ubi-DaPKC^KR8Q^::GFP / TM6.* (**D**) *w; FRT^G13^ DaPKC^k06403^ / CyO; ubi-DaPKC::GFP / TM6; w par-6::GFP hs-FLP par-6^Δ226^ FRT^9-2^.*

**Figure 3: (B)** TOP: *w; GFP::aPKC / UAS-Cdc42^CA^; hs-FLP Act5C(FRT.CD2)-Gal4 UAS-RFP / +.* BOTTOM: *w; UAS-Cdc42^CA^ / +; hs-FLP Act5C(FRT.CD2)-Gal4 UAS-RFP / ubi-DaPKC^KR8Q^::GFP.* **(C)** *UAS-cdc42-RNAi, w / w; GFP::aPKC / +; hs-FLP Act5C(FRT.CD2)-Gal4 UAS-RFP / +.*

**Figure 8:** (**A,B**) *w; lgl::GFP hs-FLP/ lgl::GFP UAS-crb; Act5C(FRT.CD2)-Gal4 UAS-RFP / +; w; lgl^S5A^::GFP UAS-Crb / +; hs-FLP Act5C(FRT.CD2)-Gal4 UAS-RFP / +.* **(C)** *w; lgl::GFP UAS-Crb / +; +/+; w; lgl::GFP UAS-Crb / αTub67C-Gal4^V2H^; αTub67C-Gal4^V37^*/ *+; w; lgl^S5A^::GFP UAS-Crb / αTub67C-Gal4^V2H^; αTub67C-Gal4^V37^/ +.* **(D)** *w; lgl::GFP; w; lgl::GFP; crb^KO^/ crb^KO^.* **(E)** *w; lgl::GFP UAS-Crb / +; hs-FLP Act5C(FRT.CD2)-Gal4 UAS-RFP / UAS-DaPKC-RNAi.* **(F)** *w; lgl::GFP UAS-Crb / UAS-par6-RNAi; hs-FLP Act5C(FRT.CD2)-Gal4 UAS-RFP / +.* **(G)** *w; lgl::GFP hs-FLP / +; Act5C(FRT.CD2)-Gal4 UAS-RFP / UAS-DaPKC*Δ*N.*

**Figure S1:** (**A**) *w / w hs-FLP; FRT^G13^ DaPKC^k06403^/ FRT^G13^ His2Av::mRFP; ubi-DaPKC::GFP/+; w / w hs-FLP; FRT^G13^ DaPKC^k06403^/ FRT^G13^ His2Av::mRFP; ubi-DaPKC^KR8Q^::GFP/+.* (**B**) *w par-6::GFP hs-FLP par-6^Δ226^ FRT^9-2^/FM7C; w; FRT^G13^ DaPKC^k06403^/CyO; ubi-aPKC::GFP/TM6.* (**C**) *w par-6::GFP hs-FLP par-6^Δ226^ FRT^9-2^/ w; lgl^KO^ FRT^40A^/ ubi-RFP^NLS^ FRT^40A^.*

**Figure S3:** (**A, B**) *w UAS-mRFP-FKBP-5’Ptas / par-6::GFP hs-FLP par-6^Δ226^ FRT^9-2^; +/+; Act5C(FRT.CD2)-UAS-RFP^NLS^/ UAS-lck-FRB::CFP*.

## Online Supplementary Material

Figure S1 shows that the polybasic PSr was required for PM targeting of *Drosophila* aPKC and both Par-6::GFP and DaPKC::GFP showed hypoxia-sensitive PM localization in embryonic epithelia. In addition, hypoxia-sensitive PM localization of Par-6::GFP was not affected by the loss of Lgl.

Figure S2 shows that in MCF7, COS7 and polarized MDCK cells PM targeting of PKCζ required co-expression of Par-6.

Figure S3 shows that acute depletion of PIP2 in Drosophila follicular cells did not strongly inhibits the PM localization of Par-6::GFP.

Figure S4 shows that Crb-intra was capable of recruiting Par-6 to PM in HEK293 cells, and that Lgl::GFP PM localization in polarized MDCK cells is resistant to the overexpression of PKCζ::RFP.

Movie S1 and S2 show the acute and reversible loss of DaPKC::GFP and Par-6::GFP from PM under hypoxia in live *Drosophila* follicular cells.

Table S1 lists the details about DNA constructs used in this report.

## SUPPLEMNTARY INFORMATION

### 1. SUPPLEMENTARY FIGURES

**Figure S1.**
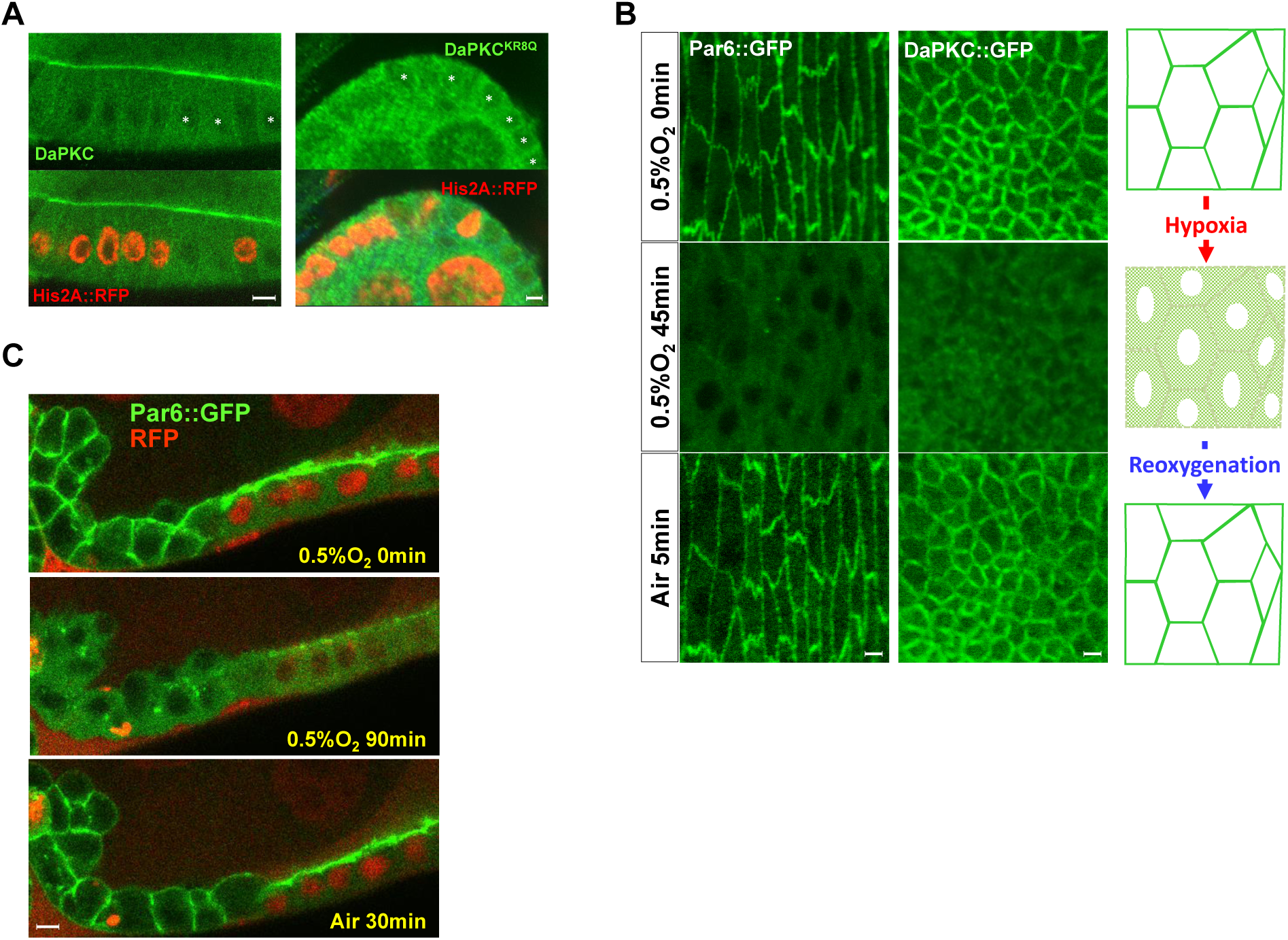
Polybasic PSr is required for PM targeting of *Drosophila* aPKC. **(A)** PM localization of wild type DaPKC::GFP and non-polybasic DaPKC^KR8Q^::GFP in *Drosophila* wild type and *DaPKC*^-/-^ mutant follicular epithelial cells. Asterisks indicate *DaPKC*^-/-^ mutant cells identified by the loss of Histon2A::RFP (His2A::RFP). Images are in cross section view. **(B)** In *Drosophila* embryonic epithelial cells, PM localization of Par-6::GFP or DaPKC::GFP was lost under hypoxia (0.5% O_2_) but recovered after post-hypoxia reoxygenation. Images are in tangential view of the apical surface of embryonic epithelia. **(C)** Par-6::GFP showed acute and reversible loss of PM targeting under hypoxia in both wild type and *lgl*^-/-^ mutant (marked by the loss of nuclear RFP) follicular epithelial cells. Note that in *lgl*^-/-^ mutant cells, Par-6 was no longer restricted to apical PM but localized to both apical and lateral PM. Images are in cross section view. Scale bars: 5µm.

**Figure S2.**
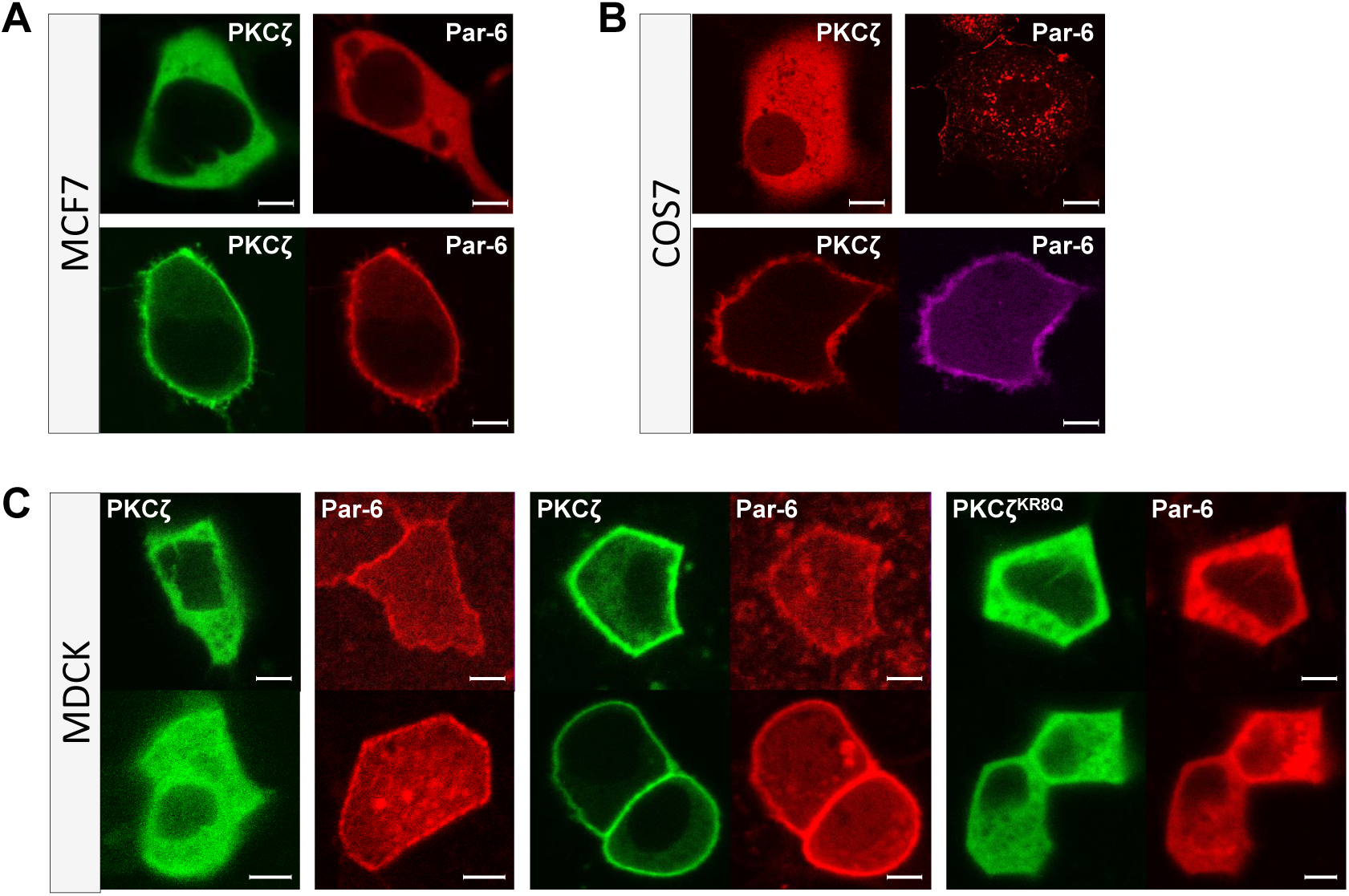
Par-6-dependent PM targeting of PKCζ in MCF7, COS7 and polarized MDCK cells. (**A-B**) Representative images showing that in MCF7 and COS7 cells, PKCζ and Par-6 were cytosolic when each expressed alone, but both became PM localized when co-expressed. (**C**) Representative images showing that in polarized MDCK cells, overexpressed PKCζ::GFP was cytosolic and Par-6::RFP was partially PM localized. Both PKCζ::GFP and Par-6::RFP were PM localized when co-expressed, whereas both PKCζ^KR8Q^::GFP and Par-6::RFP were cytosolic when co-expressed. Scale bars: 5µm.

**Figure S3.**
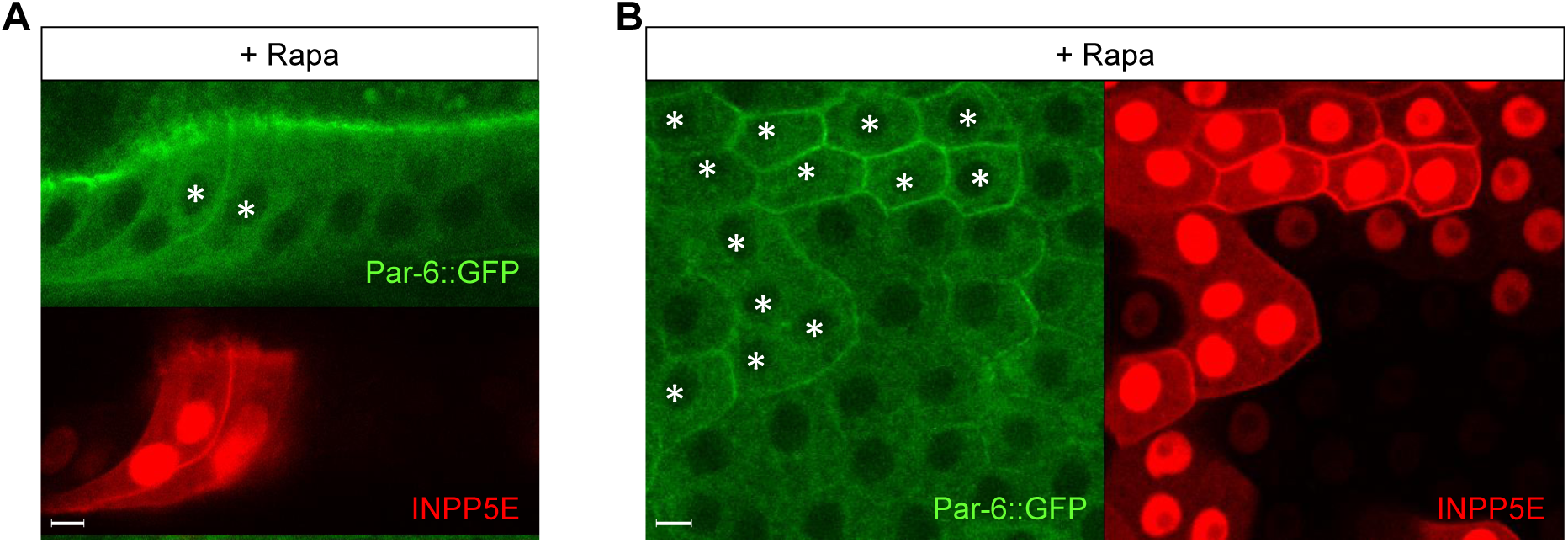
Par-6::GFP expands to basolateral PM in *Drosophila* follicular epithelial cells under the acute loss of PIP2. Cells overexpressing mRFP-FKBP-INPP5E and PM-bound Lck-FRB::CFP (not imaged) are labeled by nuclear RFP (asterisked in GFP images). Rapamycin (“rapa”) treatment induced strong PM localization of mRFP-FKBP-INPP5E, but Par-6::GFP remained largely on apical PM (**A**, cross-section view) with expansion to lateral PM (**B**, tangential view of basolateral PM). Scale bars: 5µm.

**Figure S4.**
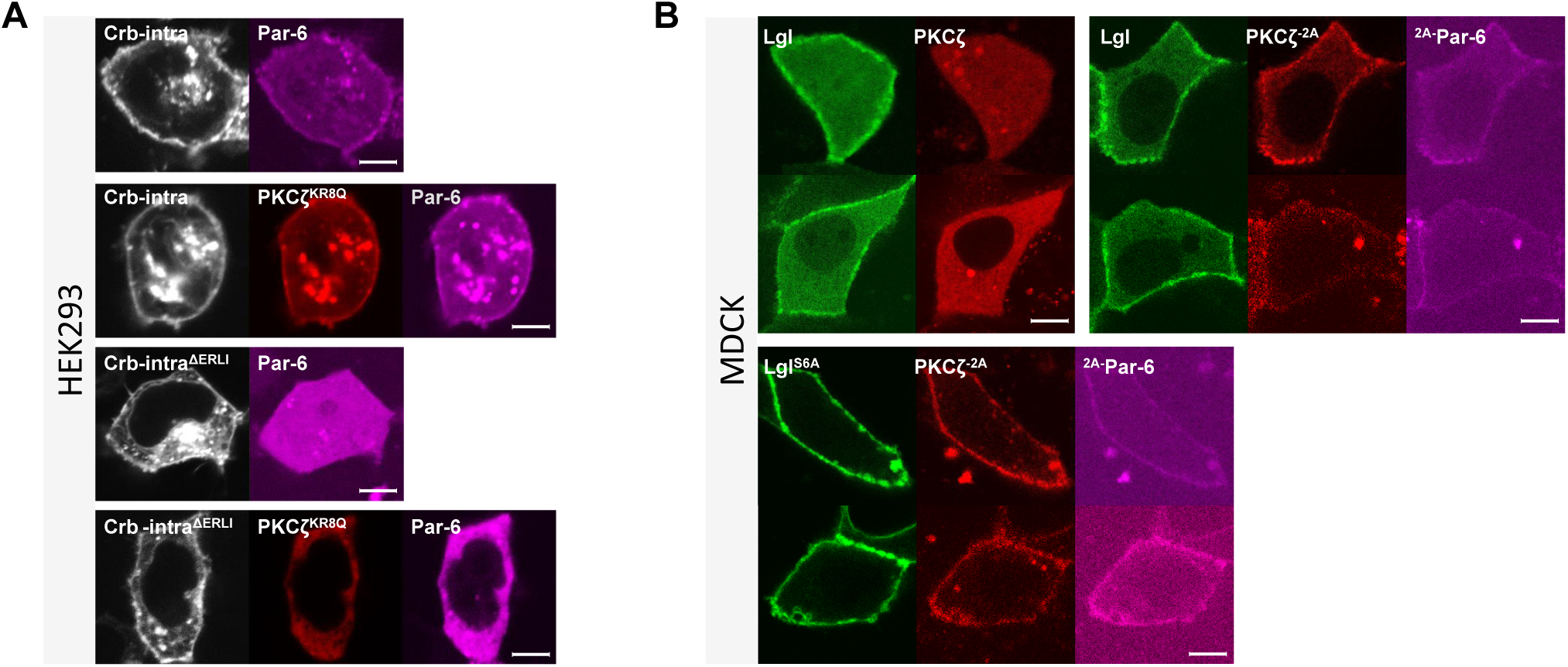
Interaction between Par-6 and Crb-intra in HEK293 cells. **(A)** Par-6::iRFP localized to PM in HEK293 cells expressing BFP::Crb-intra, but not in cells expressing BFP::Crb-intra^ΔERLI^. PKCζ^KR8Q^::RFP was PM-localized in cell expressing Par-6::iRFP and BFP::Crb-intra, but was cytosolic in cells expressing Par-6::iRFP and BFP::Crb-intra^ΔERLI^. **(B)** Lgl::GFP was predominantly PM localized in polarized MDCK cells overexpressing PKCζ::RFP or PKCζ::RFP^-2A-^Par-6::iRFP. Lgl^S6A^::GFP was PM localized in MDCK cells expressing PKCζ::RFP^--2A-^Par-6::iRFP. Scale bars: 5µm.

### 2. SUPPLEMENTARY MOVIES

**Movie S1. Acute and reversible loss of PM DaPKC::GFP under hypoxia.**

Ovaries from a 3-day old *ubi-DaPKC::GFP* female were dissected and imaged live in an environment-controlled micro chamber. Hypoxic (0.5% O_2_) gas was flashed into the chamber starting 0 minute. Normal air was flashed into chamber starting 42min for reoxygenation. Time intervals are 3 minutes during hypoxia and 10 seconds during reoxygenation.

**Movie S2. Acute and reversible loss of PM Par-6::GFP under hypoxia.**

Ovaries from a 3-day old genomically rescued *par-6::GFP par-6^Δ226^* female were dissected and imaged live similarly as samples in Movie S1. Reoxygenation starts from 33min. Time intervals are 3 minutes during hypoxia and 10 seconds during reoxygenation.

## Notes

### Competing Interest Statement

The authors have declared no competing interest.

